# The OpenPicoAmp-100k : an open-source high performance amplifier for single channel recording in planar lipid bilayers

**DOI:** 10.1101/781260

**Authors:** V Shlyonsky, D Gall

## Abstract

We propose an upgraded version of our previously designed open-source lipid bilayer amplifier. This improved amplifier is now suitable both for the use in introductory courses in biophysics and neurosciences at the undergraduate level and for scientific research. Similar to its predecessor, the OpenPicoAmp-100k is designed using the common lithographic printed circuit board fabrication process and off-the-shelf electronic components. It consists of the high-speed headstage, followed by voltage-gain amplifier with built-in 6-order Bessel filter. The amplifier has a bandwidth of 100 kHz in the presence of 100 pF input membrane capacitance and is capable of measuring ion channel current with amplitudes from sub-pA and up to ±4 nA. At the full bandwidth and with a 1 GΩ transimpedance gain, the amplifier shows 12 pA_rms_ noise with an open input and 112 pA_rms_ noise in the presence of 100 pF input capacitance, while at the 5 kHz bandwidth (typical in single-channel experiments) noise amounts to 0.45 pA_rms_ and 2.11 pA_rms_, respectively. Using an optocoupler circuit producing TTL-controlled current impulses and using 50% threshold analysis we show that at full bandwidth the amplifier has deadtimes of 3.5 µs and 5 µs at signal-to-noise ratios(SNR) of 9 and 1.7, respectively. Near 100% of true current impulses longer than 5 µs and 6.6 µs are detected at these two respective SNRs, while false event detection rate remains acceptably low. The wide bandwidth of the amplifier was confirmed in bilayer experiments with alamethicin, for which open ion channel current events shorter that 10 µs could be resolved.

## Introduction

The year 2019 marked several anniversaries in single-channel research and electrophysiology in general. Seventy years ago Kenneth Cole and George Marmont invented voltage-clamp technique [5, 12]. Same year, Hodgkin, Huxley and Katz applied this method in their work on the development of the ionic theory of action potential. In 1969, hence 50 years ago, the single ion channel gating has been demonstrated for first time; researchers from Joellen Eichner group voltage-clamped a lipid bilayer with reconstituted “excitability inducing material” obtained from *Enterobacter cloacae* [3]. Just few months later, Hladky and Haydon reported single-channel ion currents in lipid bilayers produced by gramicidine A [11]. Following these pioneering publications, single-channel ion current fluctuations have been recorded for many channel-forming molecules. Voltage-clamp technique evolved as well during these years to allow higher temporal and signal-to-noise resolution, which resulted in the development of the patch-clamp technique by Erwin Neher and Bert Sakmann in 1981 [9].

Voltage-clamp electrophysiology equipment has been commercialized since a long time. For a price counting sometimes 4-5 digits, the researcher obtains all necessary command, compensation and recording circuitry. In case of lipid bilayer studies, however, lots of electronic functionalities of these amplifiers might remain unused at all. This is true because the bilayer chamber has very low stray capacitance and because suspended bilayers have exceptionally low leakage currents and low access resistance. In addition, membrane capacitance does not have to be compensated in most instances. With the advancement in the computer and software technologies, some functions of the amplifier have been digitalized so that command voltages are not generated by the analog circuits but instead may be pre-programmed on low-cost digital-to-analog (DA) converters. Hence, the purchase of an expensive and sophisticated amplifier for bilayer studies may become unnecessary. Unfortunately, to our best knowledge, the detailed design of a low cost, discrete component, high bandwidth voltage-clamp amplifier suitable for ion channel research in lipid bilayer membranes is not available in the existing literature. Accordingly, the aim of our work was to develop an advanced version of our previous amplifier OpenPicoAmp [22] to make it suitable not only for training/teaching purposes but for electrophysiological research as well, while keeping it easily reproducible in every laboratory with minimal technical experience. The amplifier design was guided by the following considerations: 1) the setup should have wide dynamic range (from sub-pA to several nA); 2) it should have full bandwidth of 100 kHz independent of the variations of input capacitance; 3) it should be stable with large input capacitances, thus allowing experiments with classical artificial lipid bilayer membranes; 4) amplifier should have restrained number of discrete off-the-shelf components to remain easily implemented or adapted by virtually anyone. Moreover, the design using discrete electronic components should make it easy to upgrade the amplifier with the new low-noise, high-bandwidth precision integrated circuits that will be available in the future. The electronics design is licensed under a Creative Commons Attribution Share-Alike license, which allows for both personal and commercial derivative works, as long as this paper is credited and the derivative designs are released under the same license.

### Outline of the amplifier

The amplifier setup consists of two metallic enclosure cases (Figure 1A). First larger case hosts bilayer membrane chamber and smaller enclosure with input voltage divider and the headstage. Second enclosure case contains voltage-gain amplifier and 6-order Bessel filter.

**Figure 1.**
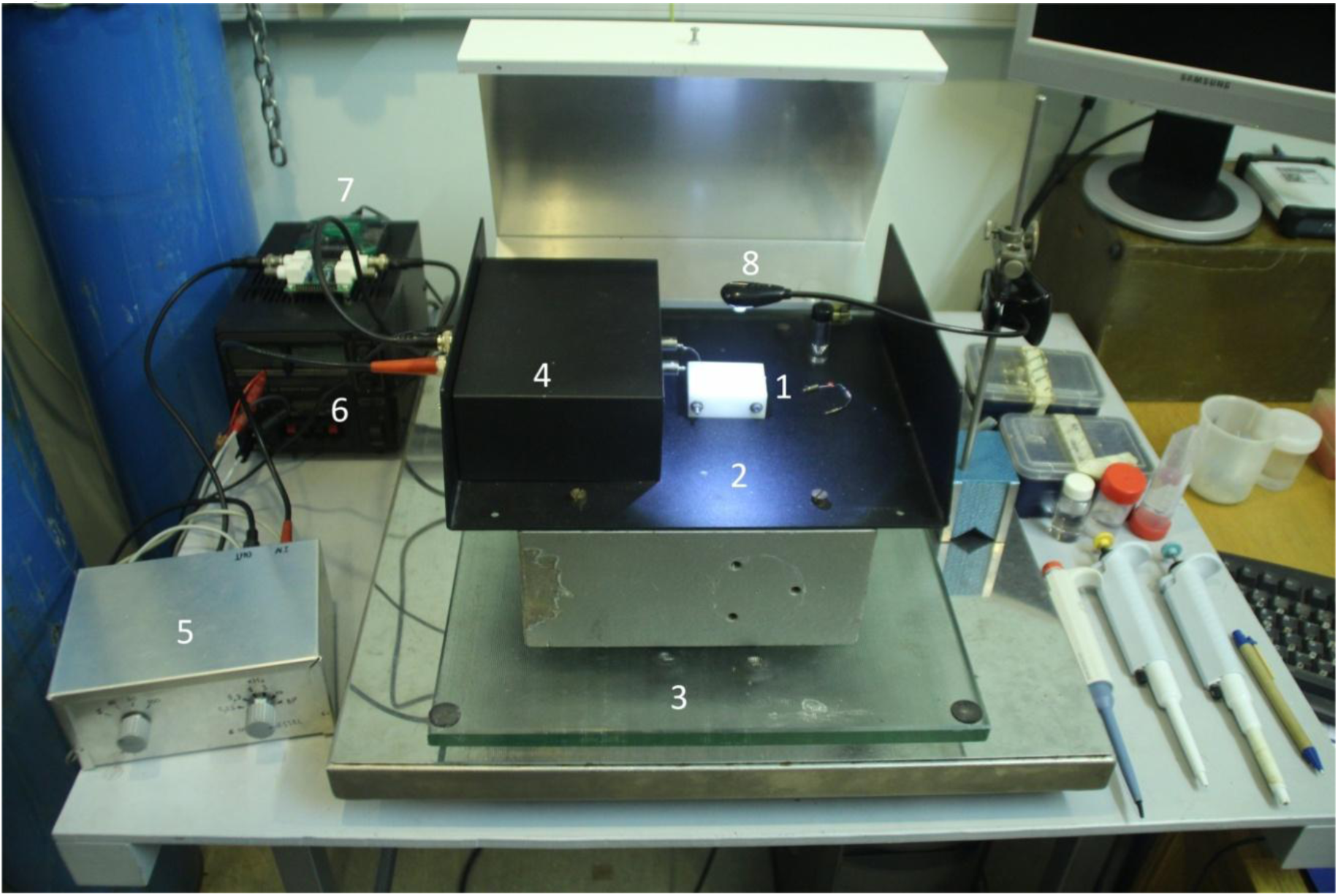
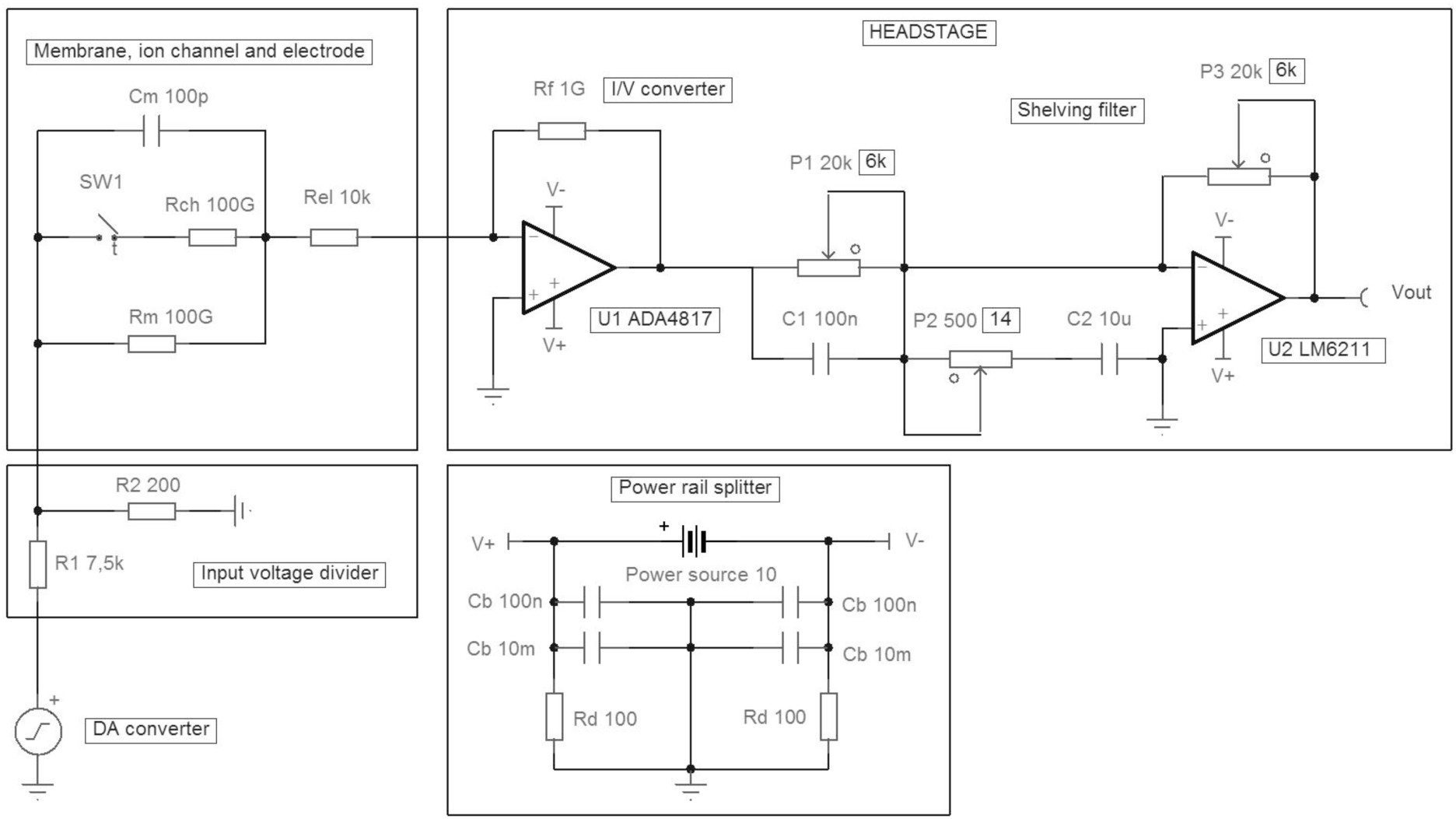
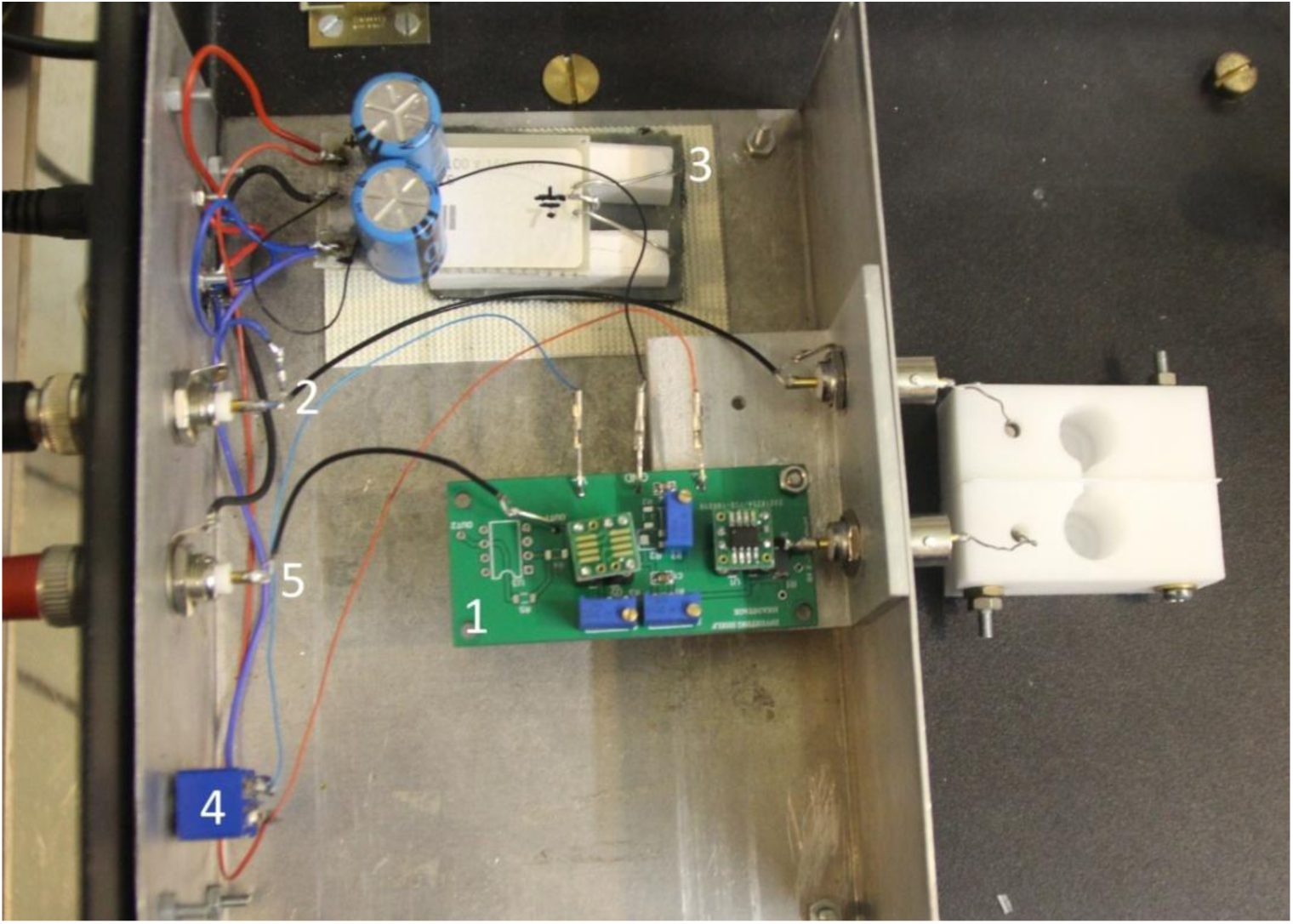
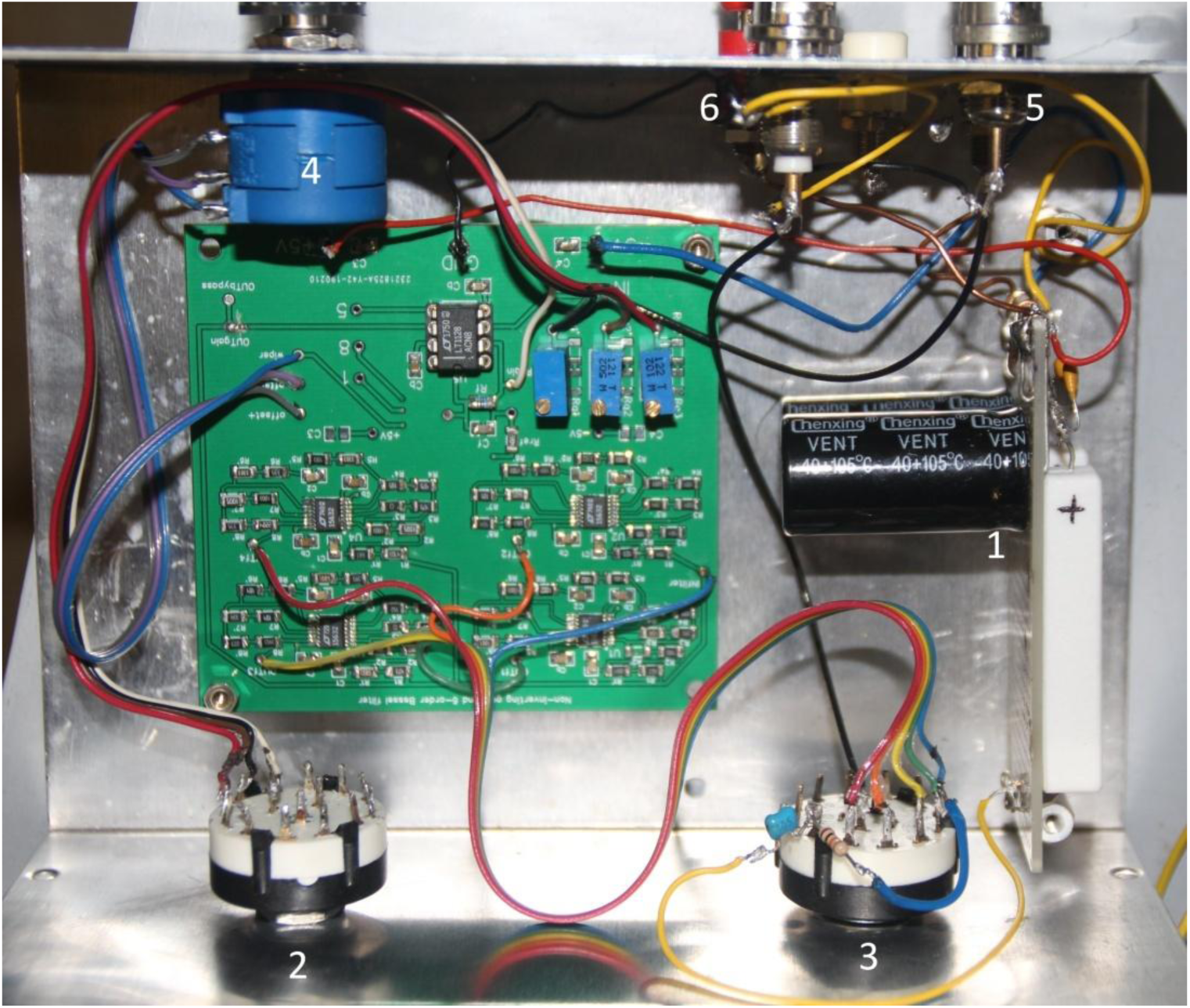
Outline of the amplifier setup. A. General view of the setup. 1-bilayer chamber, 2-bilayer chamber compartment with pivotal lid, 3-anti-vibration table, 4- headstage compartment, 5-voltage gain amplifier, 6- power source, 7-AD/DA converter, 8- light source. B. Electrical schematics of the input voltage divider, schematics of a bilayer membrane composed of membrane capacitance C_m_, membrane resistance R_m_ and ion channel R_i_, whose gating is simulated by a time-controlled switch SW1. Pair of electrodes is represented by R_el_. Power supply and headstage schematics are also shown. Bypass capacitors (10 nF and 10 µF) at the voltage supply pins of the operational amplifiers are omitted for clarity. C. Implantation of the headstage. 1- headstage PCB, 2- input voltage divider, 3- power rail splitter, 4- on/off power switch, 5- output. D. Implantation of the voltage gain amplifier. 1- power rail splitter, 2- voltage gain selector, 3- filter setting selector, 4- offset adjustment potentiometer, 5- input, 6- output.

### Power supplies and grounding

Each box is powered via isolated panel socket with a 10 V single voltage that is not referenced to ground. A low-ripple laboratory power source (3 Watts minimum) is an important requirement for a low-noise performance of the amplifier. Good results were obtained using HQ-Power PS603, PS1502A and equivalent laboratory power units. Alternatively, we have designed relatively simple and very low-noise single-voltage power supply based on BD139 transistors. Details of this power source (schematics, PCB fabrication file and the implementation) can be found in the Supplementary Information.

Dual voltage rails and virtual ground are generated in close proximity to the circuits. They are created using 100Ω (10W) resistor divider network as shown in schematics in Figure 1B. Since many high-speed and low noise precision operational amplifiers require maximum ±5V voltage supply, this value has been chosen for power alimentation of the complete setup. Hence, two voltage rail splitters are used in the amplifier, one per enclosure case. The 100Ω resistors should be matched to 0.1% to keep the voltage value of the virtual ground as close to zero Volts as possible. No additional grounding arrangement is required as long as the virtual ground and amplifier enclosures are interconnected and have low resistance link to the analog ground of the analog-to-digital converter via BNC cables.

### Command signal generation and input voltage divider

All necessary command voltages have to be generated by an external DA converter, preferably with at least 12 bits resolution and ±10V dynamic range. We suggest the MyDAQ (National Instruments) for analog OUT/IN acquisitions up to 200 kS/s or the Analog Discovery II (Digilent) for acquisitions up to 10 MS/s. Both are low-cost devices, for which academic discount is available on the top of their relatively low price. The MyDAQ may be controlled by the open-source WinEDR program (Strathclyde University), while Analog Discovery II is used with the free WaveForms program from Digilent. Alternatively, both interfaces may be controlled by LabView/Matlab with the use of corresponding Application Programming Interface bundles. Even cheaper alternatives exist: PSlab (https://pslab.io/), ISDS205B (Harbin Instrustar Electronic Technology, China) or USB-204 (Measurement Computing Corporation, USA).

The command signals are supplied to the bilayer chamber via the input voltage divider (Figures 1B and 1C). High quality MELF resistors should be used in this circuit to ensure low-noise performance. The command voltage generation by DA-converter should be scaled up accordingly. We found that voltage division at 1/30 to 1/40 is sufficient to eliminate the noise associated with DA conversion at the analog output of the instruments above. This division factor determines the maximal input voltage that can be applied to the experimental chamber; with ±10V dynamic range of the DA converters, up to ±250-330 mV can be fed to the bilayer chamber in increments of 25-33 µV. The signal is applied to the chamber compartment via panel mount bulkhead fitting BNC connector, Ag/AgCl electrode and agar bridge. This side of the bilayer membrane makes *cis* compartment. On the opposite side, the signal is fed to the current-to-voltage converter of the headstage via agar bridge, Ag/AgCl electrode and BNC connector. Since the inverting input of the operational amplifier of the I- to-V converter creates virtual zero Volts point, this side of the bilayer membrane makes *trans* compartment. No electrode offset nulling circuitry is required if the pair of electrodes is chlorinized in the bleach liquid (Chlorox, eau de Javel etc) with the pins interconnected, which ensures that both have the same potential. Details of electrode fabrication and cleaning can be found in our previous publication [22]. Briefly, measuring and reference electrodes are made from 7 cm long silver wire (Ø0.5 mm), one end of which is bent to helix and another is soldered to a golden pin. Fully chlorinized pair of electrodes should have maximum 5 kOhm of electrical resistance in 1M KCl solution.

### The headstage

The justification of the choice of the circuit topology for the fast-speed current-to-voltage conversion may be found in Supplementary Information. The schematic circuit is shown in Figure 1B and the implantation is presented in Figure 1C. Two sides of the corresponding printed circuit boards (PCB) are shown in Supplementary Figure 1. The headstage should be mounted in a separate case in a larger metallic box hosting the bilayer chamber. In order to exclude any mechanical noise, the PCB is to be firmly fixed using a screw and a nut on a metal adapter close to the solder termination of BNC connector coming from the *trans* compartment. We removed all PCB traces around the inverting input of operational amplifier to prevent the interference from PCB leakage currents. The lead from the termination of BNC connector to the amplifier input should be air-wired and should be as short as possible, ideally soldered directly.

The printed circuit boards were designed using EasyEDA Designer software (easyEDA.com) and printed on double-sided FR4 sheets by JLCPCB (Shenzhen JLC Electronics Co., Ltd.). The bottom side of the headstage PCB hosts power supply tracks with bypass capacitors, while the signal tracks are routed on the upper side of PCB. Both 0207 package radial through-hole 1 GΩ resistors and 2010 package SMD 1 GΩ thick film resistors may be used as a feedback element in the headstage with nearly identical performance. We provide additional ground connection point to decrease resistor’s stray capacitance (E-shunt). Here, thin wire may be soldered and then wrapped around the R_f_ resistor near the output side. The trimming resistors are of 3296W package; other resistors are either 1206 package in case of thick film SMD resistors or 0204 package in case of MELF resistors. The ceramic capacitors are of 0805 package. We recommend setting the trimming potentiometers to the values shown in the schematics prior to soldering to facilitate headstage tuning. If other operational amplifiers are to be used, some approximate resistor values may be found in Supplementary Information. Surface mount operational amplifiers (SOIC8 and SOP23 packages) were first soldered to SOIC-DIP8 adapters. As outlined in Supplementary Information, we choose ADA4817-1 operational amplifier as an input integrated circuit in I- to-V conversion unit for its low-noise and high-speed performance. Second integrated circuit of the headstage may be any low-noise precision operational amplifier with a bandwidth of at least 20 MHz. This minimal bandwidth is required to extend the frequency response of the headstage to 100 kHz. LM6211, ADA4637, AD8610, LT1028 and LT1128 amplifiers have shown good results. Note that the use of very high-speed amplifier here (ADA4817-1 for example) permits the bandwidth extension to the value above 300-400 kHz. The level of noise, however, becomes comparable to the voltage supply rails, so this high bandwidth is impractical. On the other hand, if only 20 kHz maximal bandwidth is required, then LT1055 or LT1056 precision operational amplifiers can be used in all parts of the amplifier. Assembled PCBs are extensively washed in isopropyl alcohol to remove any solder flux residues. The potentiometer P1 and capacitor C1 set the frequency-response of the amplifier, the potentiometer P3 sets DC gain, while potentiometer P2 together with capacitor C2 are used to prevent amplifier oscillations by adjusting and limiting the high-frequency gain peaking of the headstage. The headstage tuning is performed by applying periodic triangular voltage signal (10-30 V/s, 500 Hz) to a calibrated RC circuit (1 GΩ//100 pF in series with 10 kΩ) and observing the resulting rectangular output signal. The potentiometers P1 and P2 are re-adjusted to obtain maximally rectangular (flat and sharp) pulse. Then the 100 mV DC voltage is applied and the potentiometer P3 is adjusted to bring the output current (i.e. I-to-V converted voltage) to the value predicted by the Ohm’s law for the calibrated resistor, hence setting the exact 1 GΩ trans-resistance gain of the headstage. The calibration, i.e. measurement of true value of 1 GΩ resistor may be accomplished by placing this resistor into a voltage divider formed together with precisely measured 10 MΩ resistor. Hence the value of the resistor may be evaluated by sourcing 10 V DC voltage to the divider and measuring the output voltage with microvolt precision. No input offset voltage correction is required in the headstage if suggested precision operational amplifiers are used. Moreover, many precision operational amplifiers including those proposed for the headstage do not even provide offset nulling possibility. Eventual zero adjustment can be left to the second cascade of amplification. The use of the third operational amplifier in the headstage is optional; it may serve as an inverting unity-gain voltage follower, in case if the second cascade of amplification uses inverting topology. It adds, however, some noise; accordingly, low value resistors (<10 kΩ) should be used here to limit noise generation.

### The voltage-gain amplifier

In order to use the full dynamic range of the AD converter during the observation of very small ionic currents in pA range, further amplification of headstage-generated voltage is required. We propose voltage-gain amplifier in non-inverting topology. The implantation is shown in Figure 1D and two sides of the corresponding PCB are presented in Supplementary Figure 2. With a feedback resistor below 47 kΩ, the bandwidth of the amplifier remains much higher than the bandwidth of the headstage. This ensures that the effective bandwidth of the whole setup is close to the bandwidth of the headstage, at least at low amplification. LT1122, LT1128, LT1028, LT6018 and some other operational amplifiers showed near identical precision and low-noise performance. The offset adjustment may be performed using null pins of the operational amplifier (1, 5 or 8) and panel mounted multi-turn potentiometer according to the datasheet of the op-amp used. If the op-amp has no internal nulling possibility, we provide external offset adjustment option using 10 MΩ resistor connected to the inverting input of the amplifier (R_ref_, Supplementary Figure 2). In this case, the extreme pins of the panel mounted multi-turn potentiometer should be wired to the supply rails, i.e. “offset+” and “offset-” and the middle pin to “wiper”. If 47 kΩ feedback resistor is used, than the offset nulling up to ±25 mV (i.e. ±25 pA) may be accomplished. This is however at the expense of +0.47% gain error at unity gain. The error can be eventually corrected by re-adjusting DC gain of the headstage (see above). Accordingly, higher offset nulling range is possible with lower value of R_ref_, provided the readjustment of the DC gain in the headstage. It should be also noted that nulling range in this method is divided by the gain, so the offset adjustment via null pins of the amplifier is the preferred option.

The proposed PCB allows configuration of four different gains, which can be fixed or adjustable. Gain selection is performed by commuting resistors/potentiometers at the points Rg1’, Rg2’ and Rg3’ to the point “Rgain” using single-pole rotating switch. Unity gain is established when no resistor is connected to the “Rgain” point (Supplementary Figure 2).

Given the large bandwidth of the amplifier, the measurement of pA level ion channel currents requires efficient filtering. We propose a 6-order pseudo-Bessel filter based on LTC1563-2 integrated circuit. Two capacitors and six resistors set the cut-off frequency of the filter and four different circuits are used to generate four filter cut-off settings. The outputs of the filter (OUT1, OUT2, OUT3 and OUT4) as well as the by-pass output (OUT_bypass_) can be commuted to the amplifier BNC output using single-pole rotating switch. A two resistors in series placement is implemented on PCB to allow setting the value of each equivalent resistor to within 1-2% margins of the required theoretical value using commercially available SMD resistors. If different tolerance SMD resistors are available (1% and 5% for example) then the pre-selection of resistors may allow decreasing this error to below 0.5%. The Table 1 reports the values of resistors and the capacitors for filters with 0.3 kHz, 1 kHz, 3 kHz and 10 kHz cut-off frequencies. The provided PCB may be used to implement any other filters with cut-off frequencies between 256 Hz and 256 kHz according to the procedure described in the datasheet of LTC1563-2 circuit and with the help of free FilterCAD program from Linear Technologies (Analog Devices).

**Table 1.**
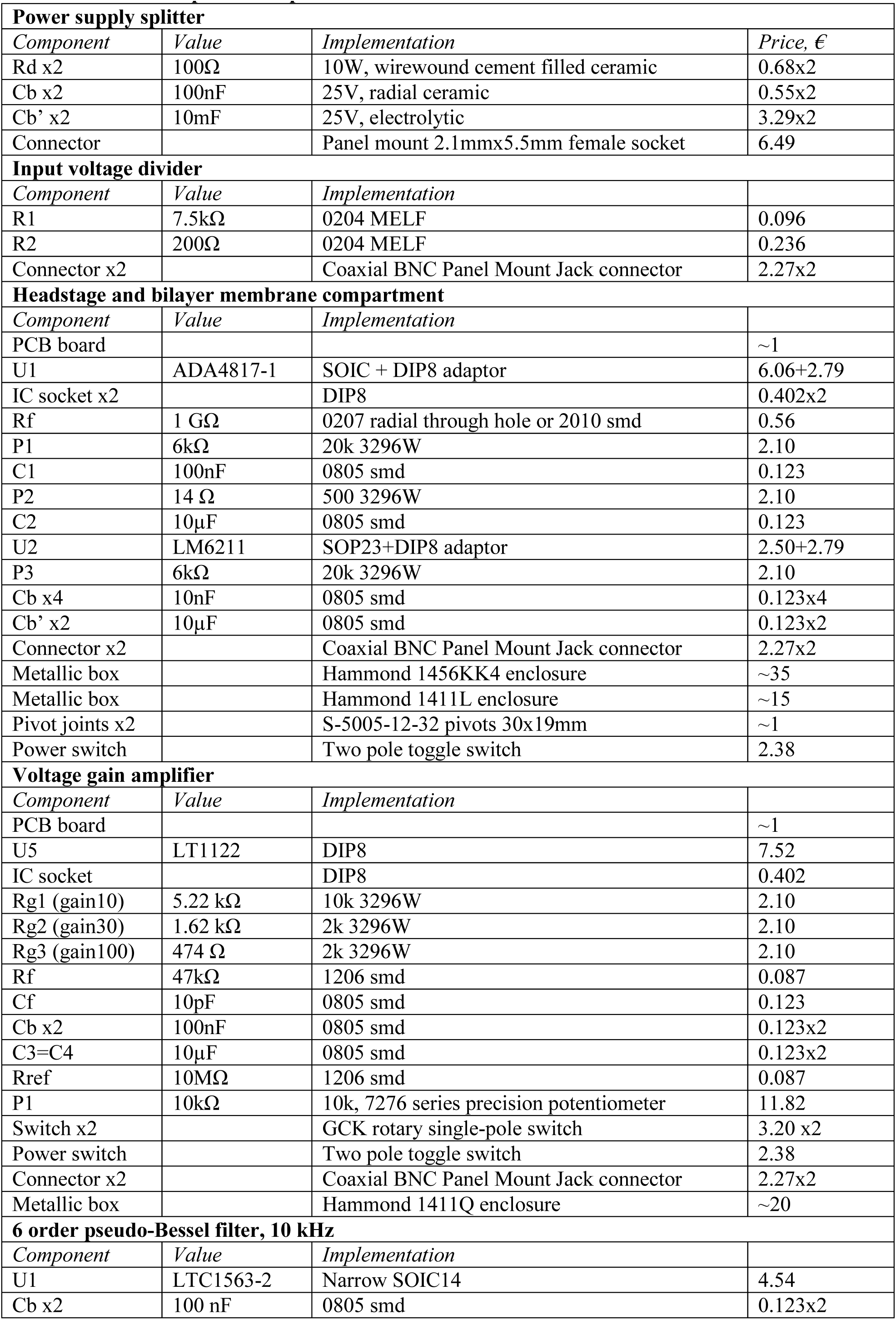
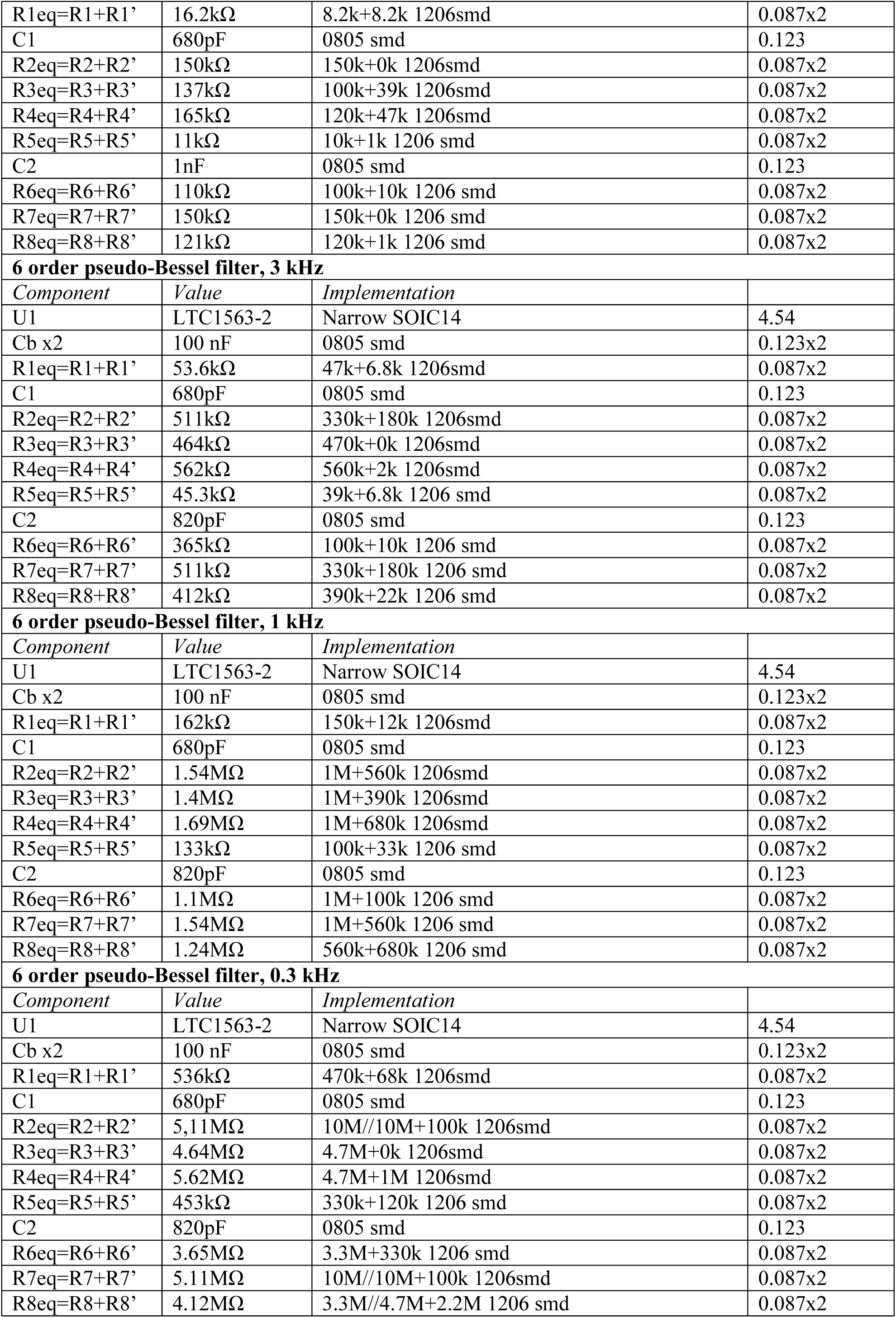
Amplifier components.

| **Power supply splitter** |  |  |  |
| --- | --- | --- | --- |
| *Component* | *Value* | *Implementation* | *Price, €* |
| Rd x2 | 100Ω | 10W, wirewound cement filled ceramic | 0.68x2 |
| Cb x2 | 100nF | 25V, radial ceramic | 0.55x2 |
| Cb' x2 | 10mF | 25V, electrolytic | 3.29x2 |
| Connector |  | Panel mount 2.1mmx5.5mm female socket | 6.49 |
| **Input voltage divider** |  |  |  |
| *Component* | *Value* | *Implementation* |  |
| R1 | 7.5kΩ | 0204 MELF | 0.096 |
| R2 | 200Ω | 0204 MELF | 0.236 |
| Connector x2 |  | Coaxial BNC Panel Mount Jack connector | 2.27x2 |
| **Headstage and bilayer membrane compartment** |  |  |  |
| *Component* | *Value* | *Implementation* |  |
| PCB board |  |  | ~1 |
| U1 | ADA4817-1 | SOIC + DIP8 adaptor | 6.06+2.79 |
| IC socket x2 |  | DIP8 | 0.402x2 |
| Rf | 1 GΩ | 0207 radial through hole or 2010 smd | 0.56 |
| P1 | 6kΩ | 20k 3296W | 2.10 |
| C1 | 100nF | 0805 smd | 0.123 |
| P2 | 14 Ω | 500 3296W | 2.10 |
| C2 | 10μF | 0805 smd | 0.123 |
| U2 | LM6211 | SOP23+DIP8 adaptor | 2.50+2.79 |
| P3 | 6kΩ | 20k 3296W | 2.10 |
| Cb x4 | 10nF | 0805 smd | 0.123x4 |
| Cb' x2 | 10μF | 0805 smd | 0.123x2 |
| Connector x2 |  | Coaxial BNC Panel Mount Jack connector | 2.27x2 |
| Metallic box |  | Hammond 1456KK4 enclosure | ~35 |
| Metallic box |  | Hammond 1411L enclosure | ~15 |
| Pivot joints x2 |  | S-5005-12-32 pivots 30x19mm | ~1 |
| Power switch |  | Two pole toggle switch | 2.38 |
| **Voltage gain amplifier** |  |  |  |
| *Component* | *Value* | *Implementation* |  |
| PCB board |  |  | ~1 |
| U5 | LT1122 | DIP8 | 7.52 |
| IC socket |  | DIP8 | 0.402 |
| Rg1 (gain10) | 5.22 kΩ | 10k 3296W | 2.10 |
| Rg2 (gain30) | 1.62 kΩ | 2k 3296W | 2.10 |
| Rg3 (gain100) | 474 Ω | 2k 3296W | 2.10 |
| Rf | 47kΩ | 1206 smd | 0.087 |
| Cf | 10pF | 0805 smd | 0.123 |
| Cb x2 | 100nF | 0805 smd | 0.123x2 |
| C3=C4 | 10μF | 0805 smd | 0.123x2 |
| Rref | 10MΩ | 1206 smd | 0.087 |
| P1 | 10kΩ | 10k, 7276 series precision potentiometer | 11.82 |
| Switch x2 |  | GCK rotary single-pole switch | 3.20 x2 |
| Power switch |  | Two pole toggle switch | 2.38 |
| Connector x2 |  | Coaxial BNC Panel Mount Jack connector | 2.27x2 |
| Metallic box |  | Hammond 1411Q enclosure | ~20 |
| **6 order pseudo-Bessel filter, 10 kHz** |  |  |  |
| *Component* | *Value* | *Implementation* |  |
| U1 | LTC1563-2 | Narrow SOIC14 | 4.54 |
| Cb x2 | 100 nF | 0805 smd | 0.123x2 |
| R1eq=R1+R1' | 16.2kΩ | 8.2k+8.2k 1206smd | 0.087x2 |
| C1 | 680pF | 0805 smd | 0.123 |
| R2eq=R2+R2' | 150kΩ | 150k+0k 1206smd | 0.087x2 |
| R3eq=R3+R3' | 137kΩ | 100k+39k 1206smd | 0.087x2 |
| R4eq=R4+R4' | 165kΩ | 120k+47k 1206smd | 0.087x2 |
| R5eq=R5+R5' | 11kΩ | 10k+1k 1206 smd | 0.087x2 |
| C2 | 1nF | 0805 smd | 0.123 |
| R6eq=R6+R6' | 110kΩ | 100k+10k 1206 smd | 0.087x2 |
| R7eq=R7+R7' | 150kΩ | 150k+0k 1206 smd | 0.087x2 |
| R8eq=R8+R8' | 121kΩ | 120k+1k 1206 smd | 0.087x2 |
| **6 order pseudo-Bessel filter, 3 kHz** |  |  |  |
| *Component* | *Value* | *Implementation* |  |
| U1 | LTC1563-2 | Narrow SOIC14 | 4.54 |
| Cb x2 | 100 nF | 0805 smd | 0.123x2 |
| R1eq=R1+R1' | 53.6kΩ | 47k+6.8k 1206smd | 0.087x2 |
| C1 | 680pF | 0805 smd | 0.123 |
| R2eq=R2+R2' | 511kΩ | 330k+180k 1206smd | 0.087x2 |
| R3eq=R3+R3' | 464kΩ | 470k+0k 1206smd | 0.087x2 |
| R4eq=R4+R4' | 562kΩ | 560k+2k 1206smd | 0.087x2 |
| R5eq=R5+R5' | 45.3kΩ | 39k+6.8k 1206 smd | 0.087x2 |
| C2 | 820pF | 0805 smd | 0.123 |
| R6eq=R6+R6' | 365kΩ | 100k+10k 1206 smd | 0.087x2 |
| R7eq=R7+R7' | 511kΩ | 330k+180k 1206 smd | 0.087x2 |
| R8eq=R8+R8' | 412kΩ | 390k+22k 1206 smd | 0.087x2 |
| **6 order pseudo-Bessel filter, 1 kHz** |  |  |  |
| *Component* | *Value* | *Implementation* |  |
| U1 | LTC1563-2 | Narrow SOIC14 | 4.54 |
| Cb x2 | 100 nF | 0805 smd | 0.123x2 |
| R1eq=R1+R1' | 162kΩ | 150k+12k 1206smd | 0.087x2 |
| C1 | 680pF | 0805 smd | 0.123 |
| R2eq=R2+R2' | 1.54MΩ | 1M+560k 1206smd | 0.087x2 |
| R3eq=R3+R3' | 1.4MΩ | 1M+390k 1206smd | 0.087x2 |
| R4eq=R4+R4' | 1.69MΩ | 1M+680k 1206smd | 0.087x2 |
| R5eq=R5+R5' | 133kΩ | 100k+33k 1206 smd | 0.087x2 |
| C2 | 820pF | 0805 smd | 0.123 |
| R6eq=R6+R6' | 1.1MΩ | 1M+100k 1206 smd | 0.087x2 |
| R7eq=R7+R7' | 1.54MΩ | 1M+560k 1206 smd | 0.087x2 |
| R8eq=R8+R8' | 1.24MΩ | 560k+680k 1206 smd | 0.087x2 |
| **6 order pseudo-Bessel filter, 0.3 kHz** |  |  |  |
| *Component* | *Value* | *Implementation* |  |
| U1 | LTC1563-2 | Narrow SOIC14 | 4.54 |
| Cb x2 | 100 nF | 0805 smd | 0.123x2 |
| R1eq=R1+R1' | 536kΩ | 470k+68k 1206smd | 0.087x2 |
| C1 | 680pF | 0805 smd | 0.123 |
| R2eq=R2+R2' | 5.11MΩ | 10M//10M+100k 1206smd | 0.087x2 |
| R3eq=R3+R3' | 4.64MΩ | 4.7M+0k 1206smd | 0.087x2 |
| R4eq=R4+R4' | 5.62MΩ | 4.7M+1M 1206smd | 0.087x2 |
| R5eq=R5+R5' | 453kΩ | 330k+120k 1206 smd | 0.087x2 |
| C2 | 820pF | 0805 smd | 0.123 |
| R6eq=R6+R6' | 3.65MΩ | 3.3M+330k 1206 smd | 0.087x2 |
| R7eq=R7+R7' | 5.11MΩ | 10M//10M+100k 1206smd | 0.087x2 |
| R8eq=R8+R8' | 4.12MΩ | 3.3M//4.7M+2.2M 1206 smd | 0.087x2 |

### Amplifier testing

The bandwidth of the complete setup was estimated from the analysis of the output rectangular signal obtained by the application of periodic triangular voltage waveform from the signal generator (GW Instek SFG-2120) to the RC test circuit [18]. This test circuit is composed of 10 kΩ resistor in series with either 100 pF or 10 pF or 1 pF capacitors to simulate different membrane capacitances. The figures 2A, 2B and 2C show the typical responses of the amplifier in the presence of these input capacitances, respectively. The signals of 10 pF and 1 pF circuits were also amplified using provided voltage-gain amplifier 10 and 100 folds, respectively, and they are shown in Figures 2D and 2E. The bandwidth (BW) of the complete setup in the presence of different membrane capacitances was estimated from the measurement of the rise time (τ_10-90_) of the sweeps and using following equation derived from the voltage relaxation equation of RC circuits [24]:

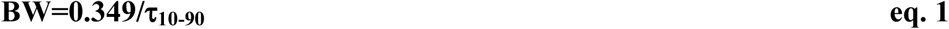

**Figure 2.**
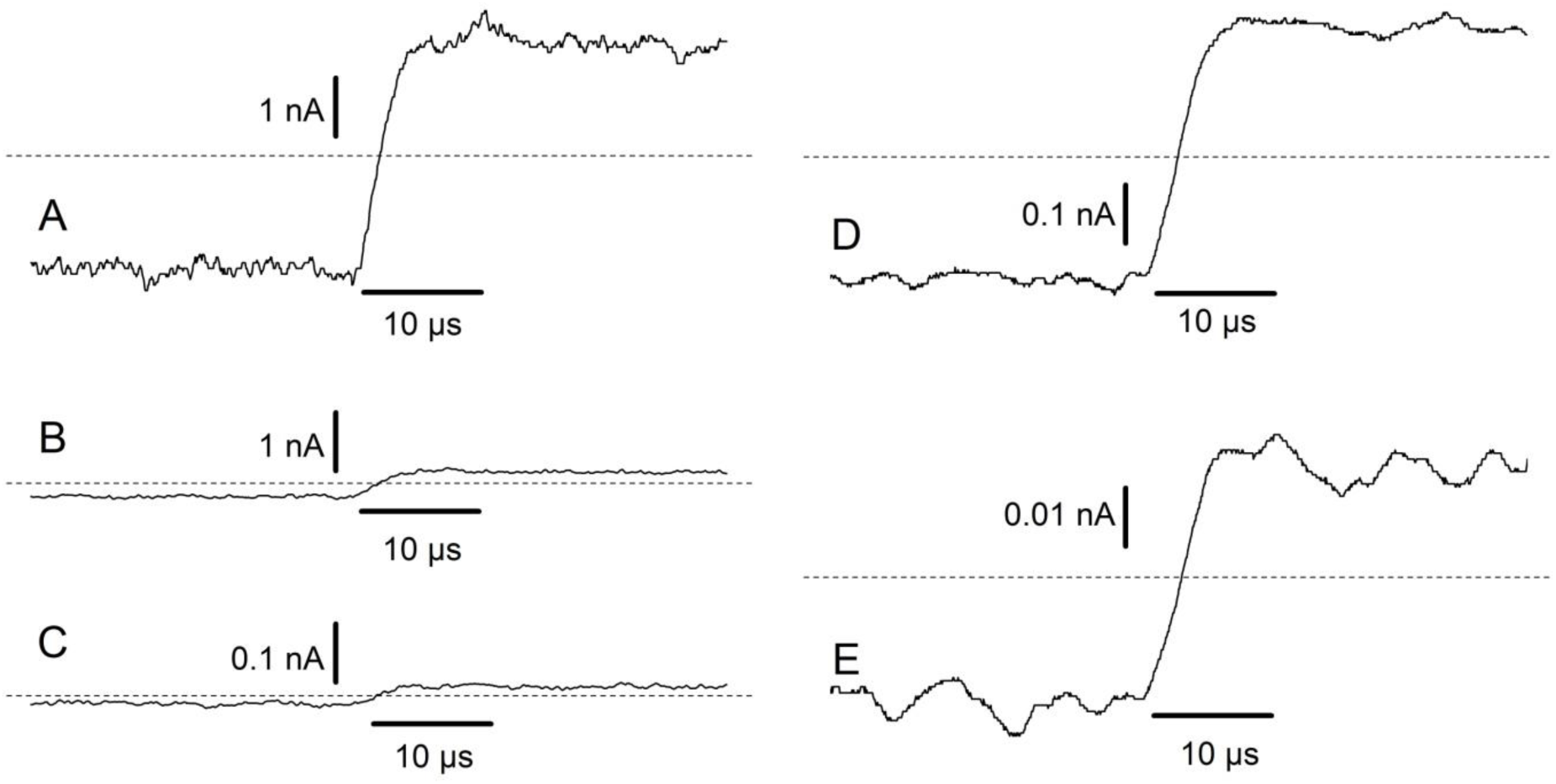
Response of the amplifier in the presence of different capacitive input loadings. Periodic triangular signal (20 V/s, 500 Hz) was applied to a test circuit composed of 10 kΩ resistor in series with 100 pF (A), 10 pF (B,C) and 1 pF (D,E). In A, C and D, recordings at the gain of 1 mV/pA are presented. In C, the signal is obtained at the gain of 10 mV/pA, while in E the gain of 100 mV/pA was used. No filtering was applied. Dotted line indicates zero current level. Only the portion of each recording is shown, which corresponds to the time point near the down peak of the input triangular signal.

The average rise-times (τ_10-90_, Mean±SD) of 50 sweeps at unity gain (1 mV/pA) were 3.37±0.62 µs, 3.48±0.42 µs and 3.51±0.36 µs for 1 pF, 10 pF and 100 pF, respectively. The respective bandwidths are estimated to be 103.6 kHz, 100.3 kHz and 99.4 kHz. These results demonstrate that the amplifier has the declared bandwidth of 100 kHz with high input capacitance.

Noise characteristics of the OpenPicoAmp in the presence of different input capacitances were measured and compared with those of the Axopatch200B using the test circuits above. To make valid comparisons we have measured the effective bandwidth of the Axopatch200B in both resistive and capacitive modes under conditions of different input capacitance loadings (test circuits above). The testing showed that the bandwidth of the Axopatch200B lags behind that of the OpenPicoAmp (last points on the Figures 3A, 3B and 3C). It should also be noted that the bandwidths of capacitive and resistive modes of Axopatch amplifier become indistinguishable at the input capacitances higher than 10 pF (close to 50 kHz, Figure 3B and 3C). With regard to noise, at low input capacitance (≤10 pF) this commercial amplifier demonstrates better noise characteristics at low frequencies in capacitive mode compared to its resistive mode and to the OpenPicoAmp (Figures 3A and 3B), while this capacitive mode has no advantage in terms of noise performance whatsoever at higher input capacitances (100 pF, Figure 3C). At the bandwidth of 5 kHz used in typical single-channel experiments, the amplifier shows 0.45 pA I_rms_ noise with an open input, which is significantly higher than 0.06 pA I_rms_ of the Axopatch 200B capacitive mode and comparable to that of its resistive mode (0.55 pA I_rms_). However, these data have little practical value, because they represent noise characteristics in ideal conditions which are never practically achieved in actual experiments. Indeed, at the 5 kHz bandwidth and in the presence of 10 pF input capacitance, the noise of the OpenPicoAmp increases to only 0.47 pA I_rms_, while the noise of Axopatch200B in capacitive mode triples in the same conditions and becomes 0.20 pA I_rms_. At the same bandwidth and with 100 pF input capacitance (a typical value for bilayer membranes used in actual experiment) noise characteristics become indistinguishable – 2.0 and 2.11 pA I_rms_ (Axopatch 200B and OpenPicoAmp, respectively).

**Figure 3.**
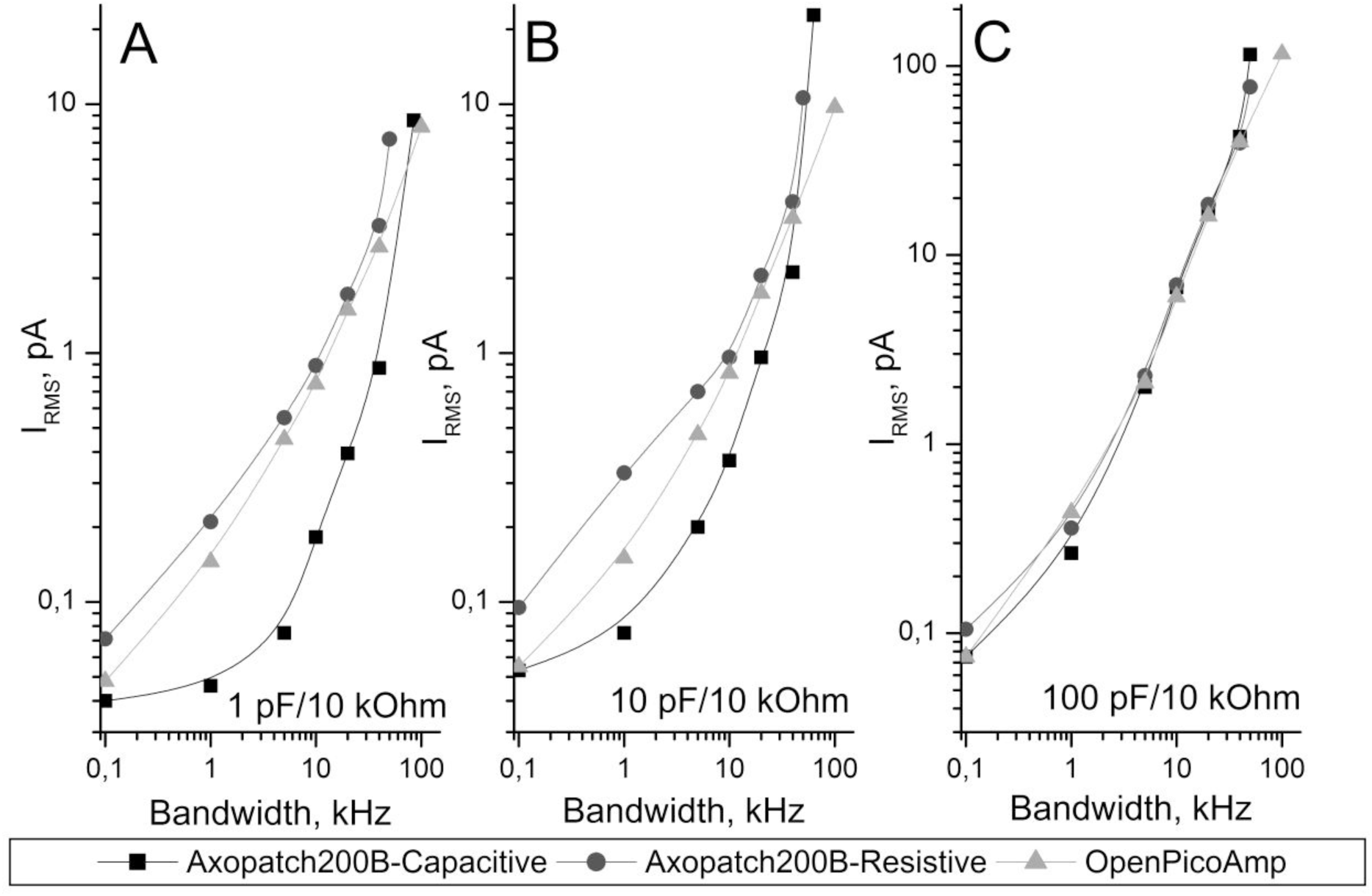
Comparison of the noise and bandwidth performance of the OpenPicoAmp and in Axopatch200B in the presence of 10 kΩ-1 pF (A), 10 kΩ-10 pF (B) and 10 kΩ-100 pF (C) input loading. RMS noise is measured either at the by-pass bandwith of the amplifiers or after filtering using 8-pole Bessel filter. Both capacitive and resistive mode maximal bandwidth (last square and circle points on the graphs) were determined from the response of the amplifier in the presence of different capacitive input loadings and under the application of periodic triangular signal. Note lower bandwidth of the Axopatch200B and near identical noise performance of all amplifiers in the presence of significant input capacitance loading.

To measure the maximal temporal resolution of the amplifier we employed a method described by Bertrand et al and by Draguhn et al [4, 8] with some modifications. The method rested on a device, which can generate TTL-controlled rectangular currents pulses in the picoampere to nanoampere range. The schematic diagram of the device is shown in Supplementary Figure 3. The current pulse is generated when a TTL-signal from a signal generator (GW Instek SFG-2120) if fed to the input of the circuit. The circuit consists of a fast-switching infra-red light-emitting diode (VSLB3940, Vishay Intertech) mounted in the feedback loop of a high-speed operational amplifier (OPA657 or ADA4817-1). The resulting light flash passes through a small aperture in a metallic shield separating two halves of a metallic box before hitting the infra-red receiver (TEFD4300F, 2 pF of distributed capacitance, Vishay Intertech). While the anode of the receiver is connected to the ground, the cathode photocurrent is sent to the input of the voltage-clamp amplifier. Note that only negative currents may be generated by this device. The amplitude of the final electrical signal can be regulated by adjusting the potentiometer and/or by positioning the infra-red receiver away from or closer to the aperture. The rise time of the VSLB3940 LED is 15 ns, while the rise time of the TEFD4300F receiver is 100 ns. These values suggest that proposed current-generating device has at least 3 MHz bandwidth with the above mentioned operational amplifiers. This is significantly higher than the 100 kHz bandwidth of the OpenPicoAmp to be tested. We can reasonably consider that the generated current impulse is highly rectangular and that it can be used for temporal resolution testing.

The Figure 4 shows typical responses of the amplifier to the current impulses of 5 µs, 10 µs and 100 µs in duration, respectively (Figures 4A, 4B, 4C). The data were acquired at 16 MHz using ISDS205B digital storage oscilloscope. For comparison, the response of the Axopatch200B amplifier in resistive mode is shown for the impulses of 20 µs and 100 µs (Figs 4D and 4E, respectively). Visual inspection of these graphs unequivocally points out again to the higher temporal resolution of the OpenPicoAmp: its 100 µs impulse is much sharper than that of the Axopatch200B. Moreover, the shape of the response of the OpenPicoAmp to 10 µs impulse is similar to the shape of the response of the Axopatch200B to 20 µs impulse again proving the bandwidth improvement (Figure 4 B and 4D, respectively). Finally, comparing the amplitudes of signals in Figures 4A and 4C suggests that the response to 5 µs current impulse preserves approximately 50% of the signal amplitude. It should be mentioned that the signal of the Axopatch200B under this condition is indistinguishable from the noise (data not shown). Unfortunately, the Axopatch200B in capacitive mode was not stable with the input signal from the optocoupler device and started to oscillate, which precluded the testing. This observation highlights increased stability of our amplifier compared to the Axopatch200B. We used 50% threshold method to estimate the current impulse duration at the level of threshold using event list generation module of WinEDR program (Strathclyde University). True signal-to-noise ratios (SNR) of 1.7 and 9 were chosen to simulate situations that are close to observing the minimal detectable signal and the optimal detectable signal, respectively. We define a minimal detectable current (I_md_) as a current with the intrinsic amplitude of 3 dB above the RMS noise and we calculate it as a ratio of the output V_rms_ and the transimpedance gain (i.e. the value of R_f_), the result being multiplied by √2 :

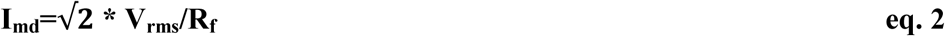

**Figure 4.**
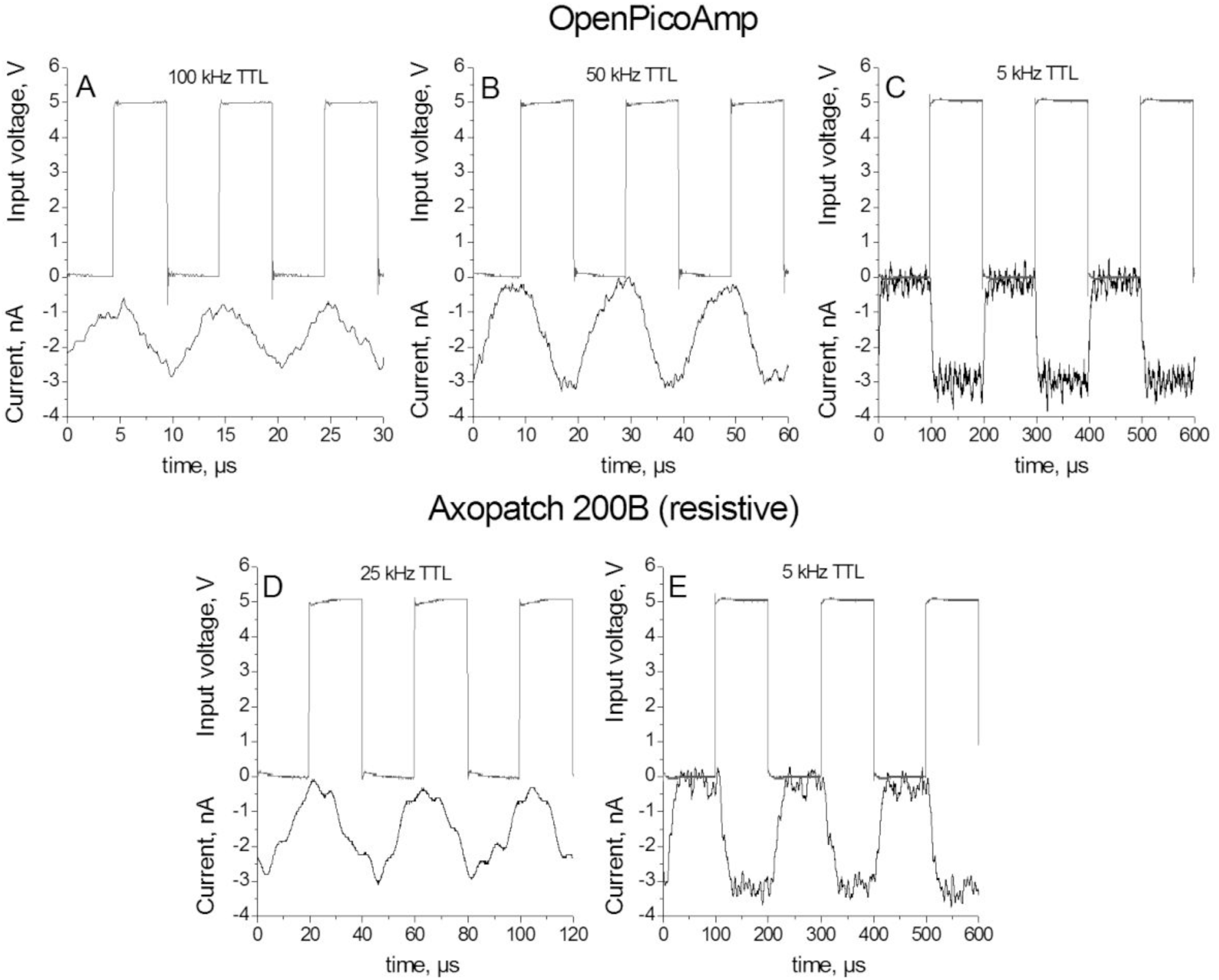
Responses of OpenPicoAmp (A,B,C) and Axopatch200B (D,E) to the current pulses with durations of 5 µs (A), 10 µs (B), 100 µs (C) and 20 µs (D), 100 µs (E). Input TTL signal is also shown for clarity (upper scale). Data were acquired at 16 MHz and presented without filtering.

In other words, minimal detectable current event with a duration corresponding to the bandwidth corner frequency has the apparent signal-to-noise ratio of 1. It is also evident that the ratio V_rms_/R_f_ is equivalent to I_rms_ at the given voltage gain. Next, optimal detectable current event at given bandwidth (I_opt_) should have the intrinsic amplitude that is 3 dB above the peak-to-peak noise, which generally corresponds to the apparent signal-to-noise ratio of 6.6 at the bandwidth corner frequency. It follows:

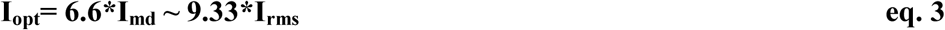

At the high SNR, only minimal post-recording filtering was used (0.8 MHz digital Gaussian). At the low SNR, more robust filtering was required to resolve the events (50 kHz digital Gaussian). Measured impulse durations were obtained from the Gaussian fitting of the event distribution histograms, one example of which at low SNR is shown in Figure 5A. These durations were plotted as a function of input impulse duration determined by the TTL frequency (Figure 6A). In accordance with the theoretical predictions [6], the measured time deviated from the linear relationship for the impulses shorter than 10 µs, which corresponds to the full rise time at 100 kHz that is the bandwidth of the OpenPicoAmp. The extrapolations of the curves to the abscissa give the values of the dead times, which were 3.5 µs and 5 µs for SNR 9 and 1.7, respectively. These dead times indicate the actual duration of a channel opening event that would give a half-amplitude response and that would pass undetectable during the post-recording analysis.

**Figure 5.**
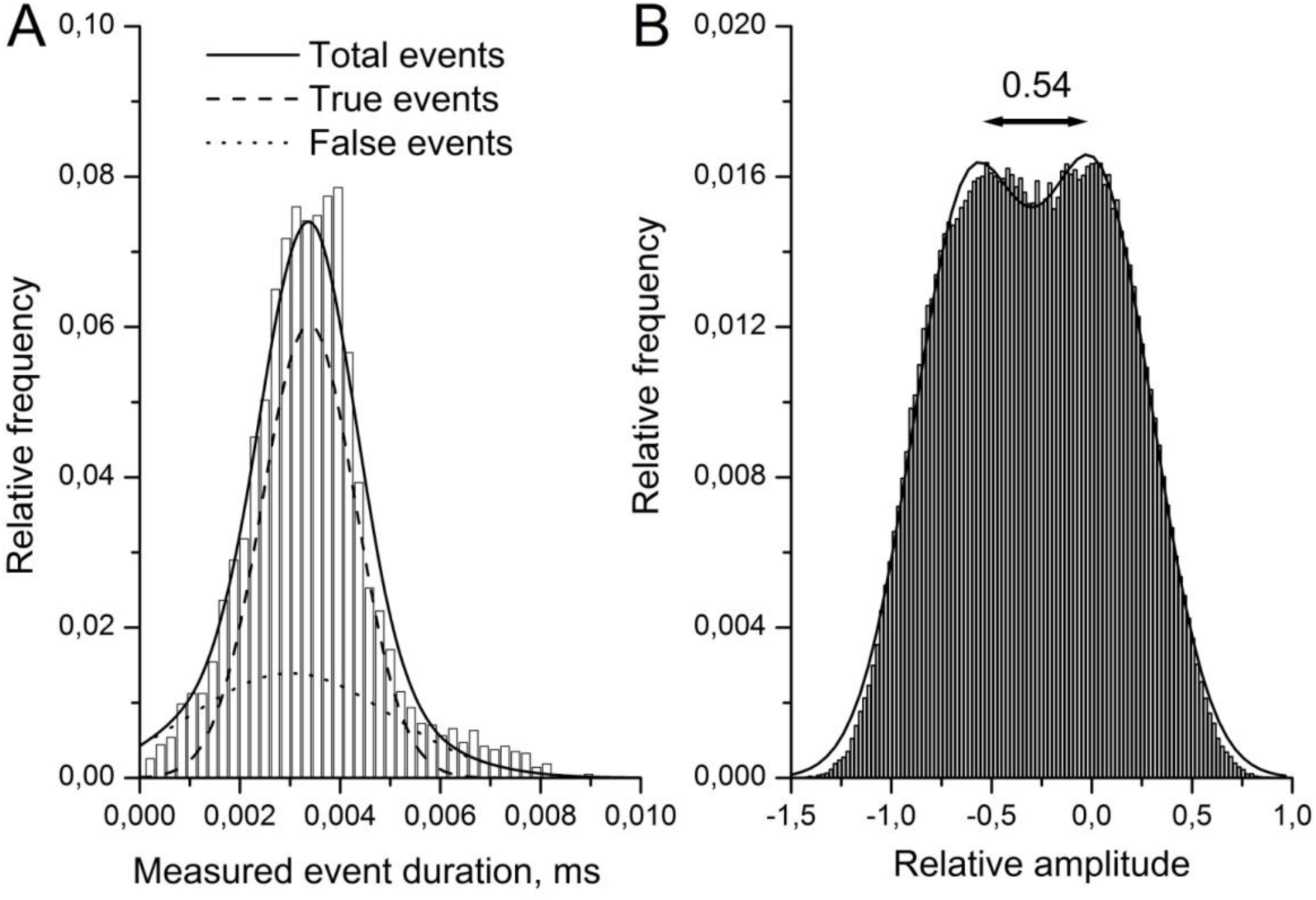
Examples of event time distributions histograms (A) and all-points histograms (B) obtained at signal-to-noise ratio of 1.7. A. Probability fraction of the events as a function of their measured duration in response to the 5 µs input current impulses. The total events distribution could be fit to the sum of two Gaussians (solid line) representing true events distribution (dashed line) and false event distribution (dotted line). B. Probability fraction of data points as a function of its current amplitude in response to the 8 µs input current impulses. Current amplitudes were normalized by the mean current amplitude observed for the 100 µs input impulse. Arrows indicate estimated relative amplitude between the two peaks.

**Figure 6.**
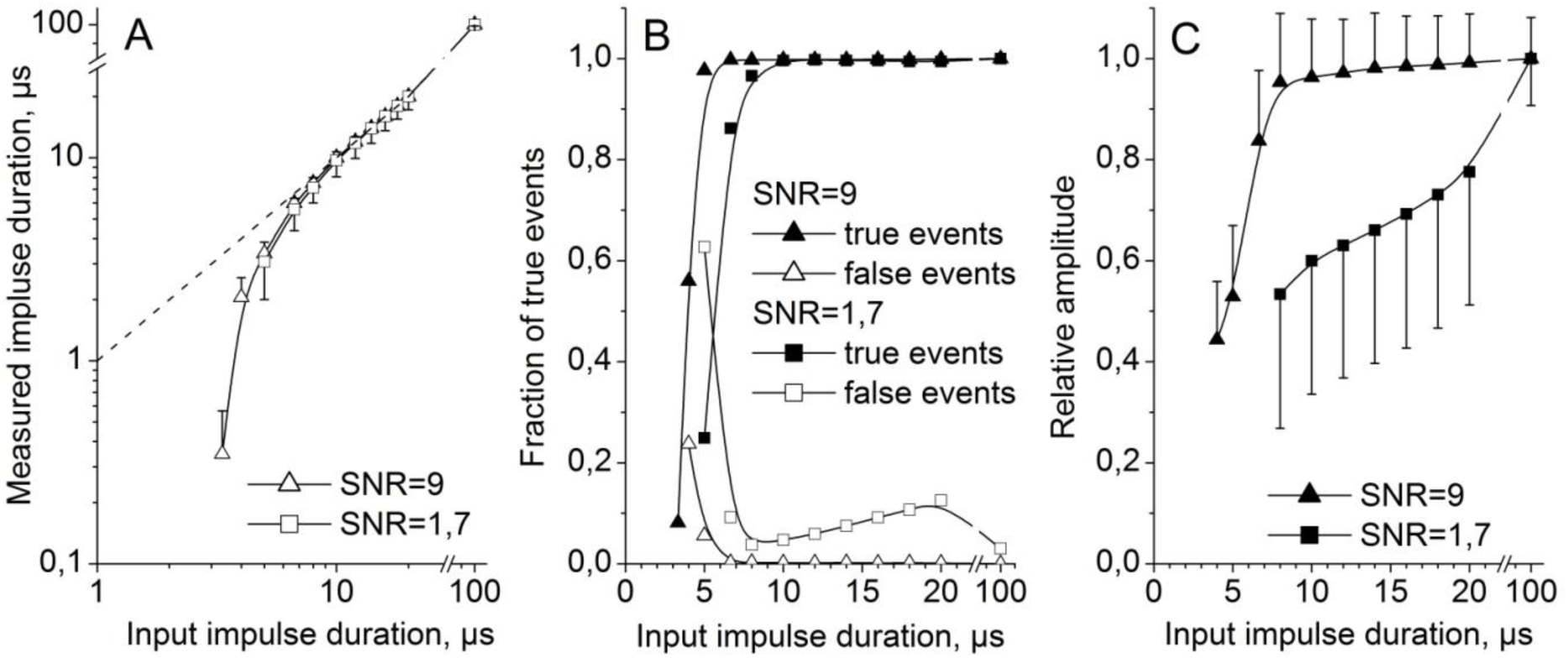
Analysis of data obtained using optocoupler device at high signal-to-noise ratio (triangles) and low signal-to-noise ratio (squares). Data were digitally filtered at 0.8 MHz at high SNR and at 50 kHz at low SNR. A. Measurement of a deadtime of the amplifier. Pulse duration at the 50% amplitude was estimated from the Gaussian fit to the measured pulse duration distribution histogram and presented as a function of input pulse duration. Dotted line indicates linear function. B. Detection of true events (inside pulse duration distribution histogram) and false events (outside pulse duration distribution histogram) expressed as a fraction of the input pulse number and as a function input pulse duration.

Given the fact that TTL frequency determined the number of true current events sent to the amplifier, we were able to calculate the fraction of input events detected. At the same time, events that fell outside of Gaussian fit of the event time distribution histograms represent the false signal and their numbers were also estimated. The examples of true and false event distribution at low SNR are shown in Figure 5A, at high SNR these distributions give sharp peaks and they are not shown. The true and false events numbers detected were normalized by the number of input events under each condition and the results are presented in Figure 6B. They show that more than 80% of true events with the durations of 5 µs and 6.6 µs are detected at the SNR of 9 and 1.7, respectively. Longer events are detectable at 100%. The false event number represents approximately 10% of true events at the SNR=1.7 and approaches 0% at higher SNR. These numbers are overall in agreement with the theoretical considerations [6].

We then next estimated the effect of post-recording filtering on the amplitude of the current impulses. All-point histograms were constructed and were fit to the sum of Gaussian functions. The representative histogram obtained at low SNR is shown in Figure 5B. The results of the analysis are presented in Figure 6C. As expected, 0.8 MHz filtering at SNR=9 did not have any impact on the current amplitudes, that remained flat in the pass-band of the amplifier. On the contrary, 50 kHz post-filtering at the SNR=1.7 expectedly decreased the amplitudes of current events shorter than 20 µs, which however remained above the 50% threshold value that allowed their subsequent resolution and detection down to 8 µs. Below this value, the peaks on the all-point distribution histograms were not resolvable. It is clear that in case of a post-filtering below the amplifier and/or filter bandwidth used during recording, the current amplitude value of events with frequencies above the filter cut-off has to be corrected. In conclusion, the OpenPicoAmp-100k permits the resolution and detection of the current events as short as 10 µs even at low signal-to-noise ratios. It should be further noted that the extrapolation of data to -3dB point on the curve for SNR=9 (Figure 6C, triangles) gives a time point value of 5.97 µs that corresponds to 167.5 kHz bandwidth frequency. This higher apparent bandwidth is achieved due to the omission of 10 kΩ resistor in series with infrared receiver capacitance. This observation highlights once again the importance of the bandwidth testing under realistic model settings closely resembling actual conditions in bilayer experiments.

We have accordingly performed alamethicin ion channel recordings in planar lipid bilayer membranes. Lipid bilayers were formed by an air bubble wicking technique [7, 22] over an aperture of 20-30 µm in diameter made in 10 µm thick PTFE film (Sigma-Aldrich, Overijse, Belgium). This aperture has been produced using the tip of a glass capillary pulled for patch-clamp studies and chopped off to the desired diameter under microscope and with the aid of a micromanipulator. The film was then positioned on a greased 1 mm thick PTFE sheet with the aperture situating in the center of a predrilled orifice of 1 mm in diameter. The bilayer membranes were formed by the air bubble formed at the tip of a micropipette pre-wetted in membrane-forming solution (10 mg/ml azolectin in n-octane, both from Sigma-Aldrich, Overijse, Belgium). Air bubble wicking is performed on the side of the film. Figure 7 shows an example of recording that was obtained in 2 M KCl symmetrical solutions in the presence of 1 pM of alamethicin and under 200 mV of applied voltage. Data were acquired at 16 MHz and were presented either unfiltered, filtered using 50 kHz digital Gaussian filter or denoised using discrete wavelet transform. The total capacitance estimated from the capacitive current amplitude was 25 pF. The current amplitudes of the first and the second apparent open levels were determined from the all-point histograms and were 112 pA and 443 pA, respectively. These correspond to the zero-voltage conductances of 560 pS and 2217 pS, respectively. The obtained unitary current conductances represent most likely the second and the third open states of alamethicin [13, 15] with the first state, which is below 50 pS in 2 M KCl, being not resolvable at the 100 kHz bandwidth. In this regard it should be mentioned that RMS noise of the membrane with fully closed channels was 29.5 pA. This implies a SNR of approximately 0.3 for this smallest open state (i.e. 50 pS) under our experimental conditions that is obviously below the limit of detection. One could nevertheless observe the difference between the noise of the membrane with fully closed channel and the noise of this smallest open state at the end of recording in Figure 7. The inserts in the graph demonstrate fully resolved event marked by an asterisk. This is a transition between 50 pS, 560 pS and 2217 pS and back to 50 pS conductance levels in approximately 10-15 µs. Another resolved event of a transition between 560 pS and 2217 pS and back to 560 pS levels in about 8 µs is marked by a double asterisk. The decrease of the amplitude of this short event after filtering using 50 kHz digital Gaussian filter is clearly noticeable. For comparison, we also show the same trace that has been denoised using SURE (Stein Unbiased Estimate of Risk) thresholding based on high-level wavelet transform approach (symlet2, level 9). This method permits achieving similar increase in SNR as with 50 kHz digital filter while it preserves the amplitudes of the fastest events. Several other non-resolved current events could be observed on the recording, which confirms nanosecond scale gating of alamethicin channel [15]. Overall, the presented results demonstrate the guaranteed minimal 10 µs temporal resolution of the proposed amplifier.

**Figure 7.**
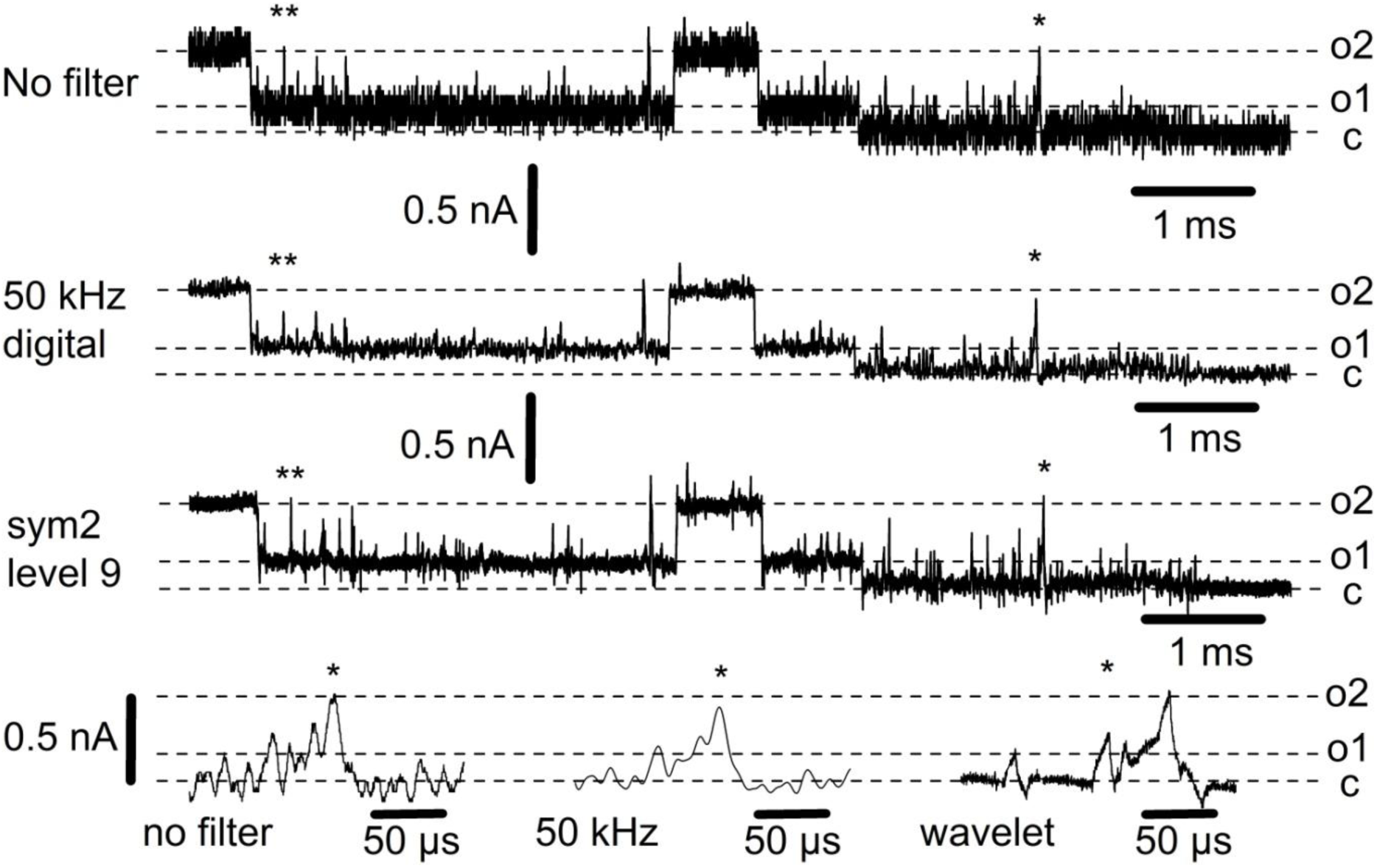
Alamethicin ion channel recording in lipid bilayer. The recording was obtained from a bilayer membrane with capacitance of 25 pF that had 29.5 pA RMS noise in 2 M KCL symmetrical solutions and under +200 mV of applied voltage. Traces present non-filtered data, data digitally filtered at 50 kHz and data obtained by a symlet-2 SURE transformation (level 9) with hard threshold rule and level-dependent noise. Time scale expansions show a brief fully resolved event marked by asterisk. Double asterisk indicates another fully resolved event of 8 µs in duration. Note the preservation of the amplitude of all fast events after wavelet transformation.

## Discussion

We describe the design and the performance of the OpenPicoAmp-100k, an open-source lipid bilayer membrane amplifier suitable for the use in introductory courses in biophysics and neurosciences at the undergraduate level as well as for the research on the high-bandwidth measurements of ion channel and nano-pore currents. This work is the outcome of the analysis and testing of the ideas that can be found in the literature (see Supplementary Information) combined with the selection of high-performance integrated circuits available in retail market at the moment. Different validation steps using the described test-circuits show that the amplifier produces low-noise, predictable and reproducible results, while actual bilayer experiments yielded expected outcomes. The previous version of the amplifier has been already used for teaching purposes [22]. The amplifier gives undergraduate students unique opportunity of hands-on approach to the understanding the basic and advanced concepts of membrane electrophysiology down to a single molecule level. The wideband version of the amplifier will extend the number of experiments that students can perform to understand the biophysical properties of ion channels in lipid membranes. The OpenPicoAmp-100k performed equally well as the benchmark amplifier Axopatch200B under real experimental conditions, i.e. in the presence of significant input capacitance (see for discussion [24]). Accordingly, the outcome of our work provides a quality research tool for investigators with lower budgets. The data presented in Figure 3 can be used to calculate minimal detectable signals at given bandwidth and in the presence of different membrane capacitances using equation 2. For example, after analog filtering at 100 Hz, 0.078 pA and 0.106 pA current events of 10 ms in duration can be resolved in membranes having 10 pF and 100 pF capacitances, respectively.

The cost of electronic components used in the described amplifier is estimated to be approximately 125 euros based on retail prices (Table 1) and the overall cost of amplifier implementation is below 250 euros that includes the benchtop low ripple power supply and metallic enclosures cases, one of which serves as a Faraday cage for bilayer chamber. Moreover, these enclosures cases can be self-made and the design of low-noise power supply for electrophysiology amplifier is also available [16, Supplementary Information]. The overall cost does not include the AD/DA interface that should be purchased separately. The vibration isolation table constructed using marble slab positioned on tennis balls proved to be sufficiently efficient in damping mechanical vibrations. Anti-vibration table consisting of two top cascaded tennis ball–suspended plates provides further improvement in vibration dumping characteristics.

The electrophysiology continues to benefit from the advances in electronic measurement systems and physiological techniques. Our work demonstrates an enormous potential for improving today’s electrophysiological equipment, since we have achieved a technological level comparable to that of state-of-the-art amplifiers with just in-house resources. In addition, we highlight the importance of an amplifier’s noise and bandwidth characterization under realistic conditions, i.e. with some resistive and capacitive input loading and not with the open input.

Further improvements of voltage-clamp amplifiers will certainly concern both bandwidth extension and noise reduction. It is generally accepted that variable high-frequency voltage-clamp current noise is a sum two major components [23-25]. First is produced by the thermal voltage noise of the access resistance in series with membrane capacitance. In many instances of bilayer experiments the contribution of this component is minor. Feedback element of the current-to-voltage converter is another source of thermal noise. In Axopatch200B this noise component is attenuated by cooling the headstage to -15°C. However, Figure 3 suggests that active cooling would significantly improve noise characteristics of the OpenPicoAmp only in the presence of input capacitance equal to 10 pF and below, which is generally difficult to achieve in real bilayer membrane experiments. For this particular reason, active cooling has not been tested in this version of the amplifier, but certainly will be considered in eventual upgrades.

Second current noise component is produced by interaction of the headstage input voltage noise with the overall input capacitance, i.e. from the membrane, stray and from that of the amplifier itself. With regard to this noise component, the development of ultra low-noise low input capacitance operational amplifiers will certainly help achieving better noise characteristics. On the other hand, bilayer lipid membrane minimization that incurs lower capacitances would also minimize noise and would extend the useful part of the available amplifier bandwidth [13, 25]. In our practice, punching thin 10 µm Teflon films with a glass patch pipette may produce extremely regular apertures of 20-30 µm in diameter. If these films are then sandwiched with a thicker Teflon sheet that minimizes stray capacitance of the film, characteristic thinning of the bilayer membrane may be electronically observed even with the painting method of bilayer formation. The capacitance of these bilayers can be as low as 20-30 pF and this greatly attenuates the overall noise, which is demonstrated in our report. To the best of our knowledge, the results shown for alamethicin channel in Figure 7 are the fastest recordings ever reported using classical bilayer membrane and discrete component amplifier as opposed to the complicated microbilayer membrane fabrication and customized integrated circuit amplifiers [2, 10, 15 and Supplementary Information]. Further reading on the suggestions and practical experimental approaches for noise optimization in high bandwidth ion channel recordings can be found in the literature [10, 13, 17, 19, 25, 26]. At last, special post-recording denoising and current idealization techniques also deserve attention when working with high bandwidth ion channel data. These techniques include discrete wavelet transforms such as demonstrated in Figure 7 [see also 1, 20, 21], idealizations based on hidden Markov model with Baum-Welch algorithms of parameters estimation [18] or model-free algorithms [14].

## Supporting information

EasyEDA source file for the gain amplifier, headstage and power supply

Gerber PCB fabrication files for the gain amplifier, headstage and power supply

## Acknowledgements

We gratefully acknowledge Freddy Dupuis for his technical help in the design of the rail splitter and Prof. Fabrice Homblé for the critical reading of the manuscript. This project has received support from the Fonds d’Encouragement à l’Enseignement of the Université libre de Bruxelles. The funders had no role in study design, data collection and analysis, decision to publish, or preparation of the manuscript.

