## Supplemental information for "The OpenPicoAmp-100k : an open-source high performance amplifier for single channel recording in planar lipid bilayers"

### Supplementary Information

#### 1. Supplementary Figures:

Figure 1. Two sides of the headstage PCB.

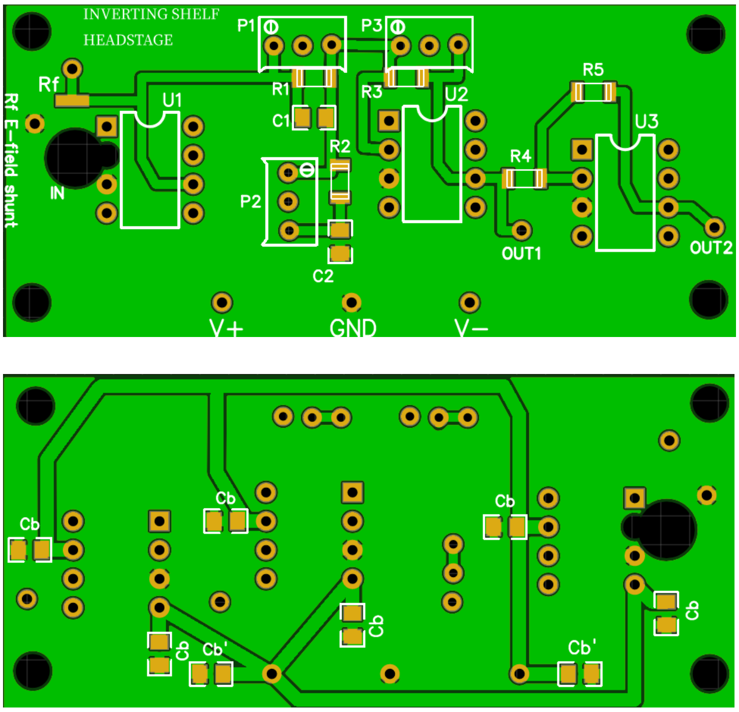


Figure 2. Two sides of the voltage-gain amplifier and 6-order pseudo-Bessel filter PCB.

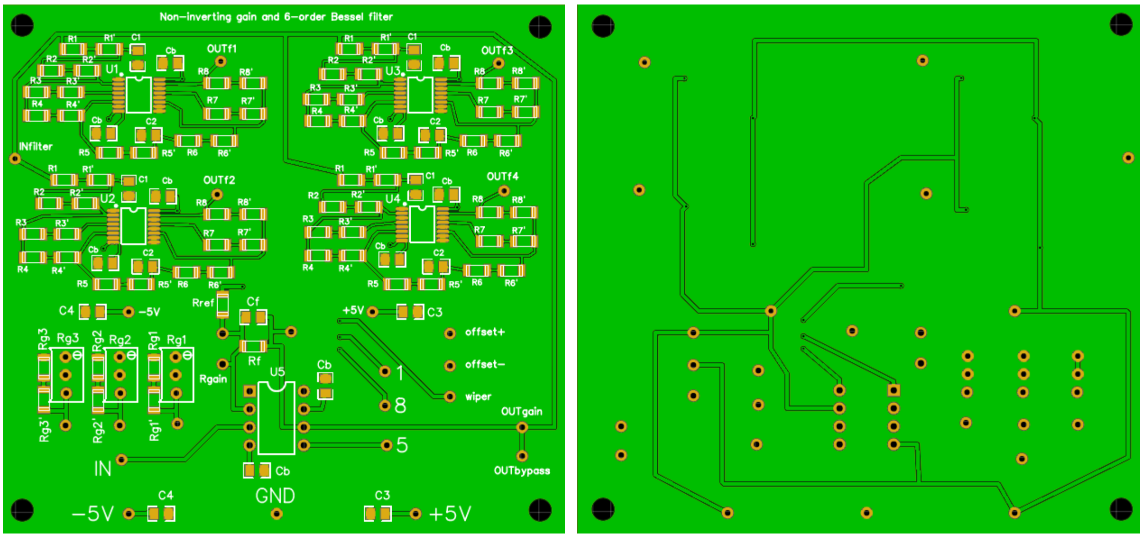

Figure 3. Electrical schematics of a current pulse generating optocoupler circuit. Light-emitting diode VSLB3940 is separated from infra-red receiver TEFD4300F by a metallic shield, in which small aperture was drilled, that is not shown.

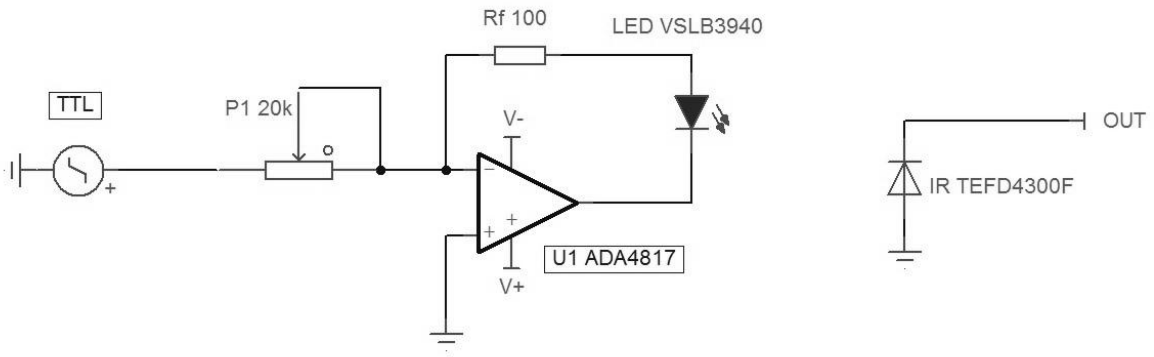


### 2. Low-noise power supply

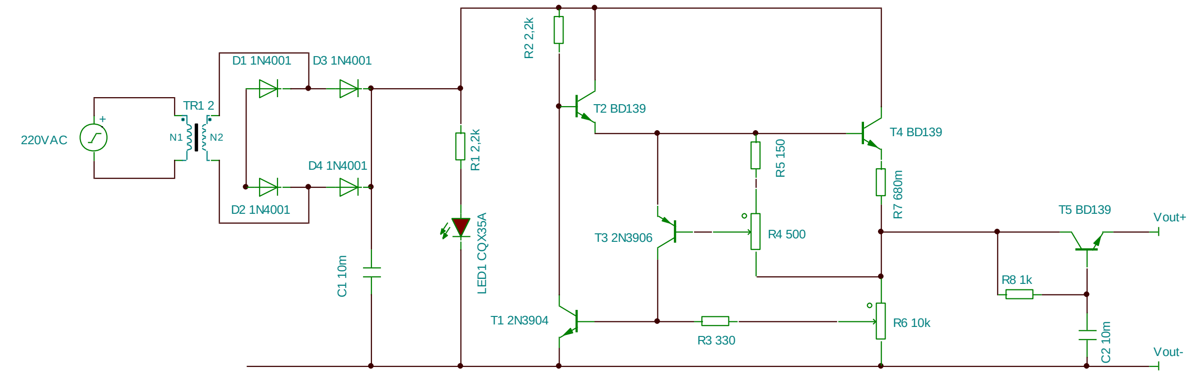


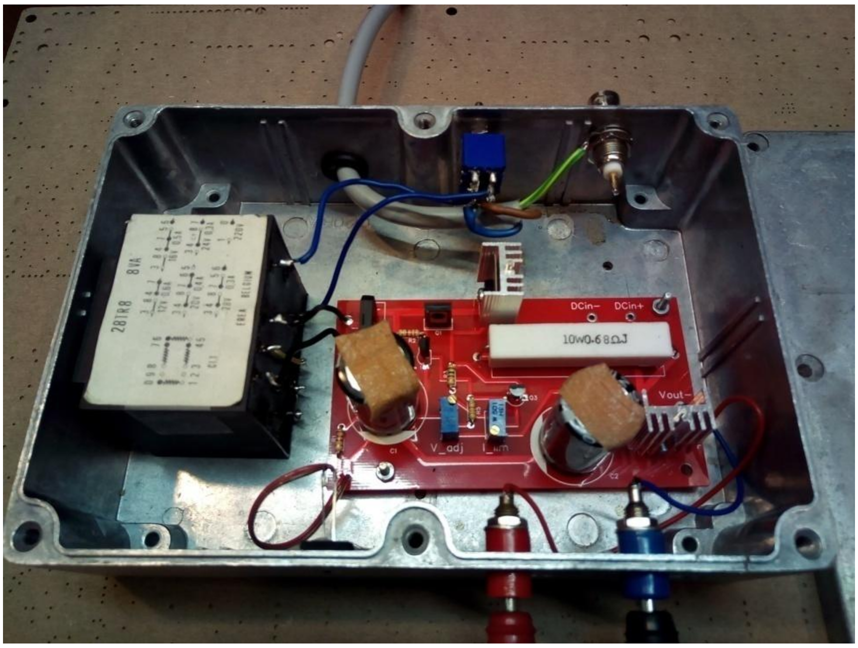

Electrical schema and actual implementation of the low-noise power source. Resistor R6 sets the output voltage (up to 24V DC) and R4 serves as a short-circuit protector by limiting the output current. Setting R4 to 50% of its value limits the output current to 250 mA that is more than sufficient for the amplifier functioning. Transistors T4 and T5 do not have to be fitted with heat-sinks, since current consumption is estimated to be maximum 150 mA. DCin+ and DCin- may be used to clean the ripple of the available DC power supply, in this case only R8, T5 and C2 are implicated. Note that the output capacitance multiplier circuit formed by R8, C2 and T5 has a time constant of 10 seconds so that it takes almost 1 minute for the output voltage to reach its adjusted value.

**Table 1. Power supply components**

| *Component* | *Value* | *Implementation* | *Price, €* |
|---|---|---|---|
| AC transformer | 220in-24out | 28TR8 8Watt transformer | ~10 |
| Diode bridge | | RS201 | 0.20 |
| C1=C2 | 10mF | 25V, electrolytic | 3.29x2 |
| R1=R2 | 2.2 kΩ | 0.25W, axial 0.3 | 0.10x2 |
| LED1 | | CQX35A red LED | 0.10 |
| T2=T4=T5 | | BD139 NPN transistor | 0.54x3 |
| T1 | | 2N3904 NPN transistor | 0.40 |
| T3 | | 2N3906 PNP transistor | 0.40 |
| R3 | 330Ω | 0.25W, axial 0.3 | 0.10 |
| R4 | 250Ω | 500 Ω 3296W potentiometer | 2.10 |
| R5 | 150Ω | 0.25W, axial 0.3 | 0.68 |
| R6 | variable | 10kΩ 3296W potentiometer | 2.10 |
| R7 | 0.68Ω | 10W, wirewound cement filled ceramic | 0.68 |
| R8 | 1kΩ | 0.25W, axial 0.3 | 0.10 |
| Power switch | | Two pole toggle switch | 2.38 |
| Connector x2 | | Panel mount « banana » female socket | 0.49x2 |
| AC Power cord | | | |
| Enclosure | | Hammond 1411Q enclosure | ~15 |
| **Total** | | | **~43** |

### 3. PCB soldering and cleaning

There are no particularly heat sensitive components in the amplifier, even suggested surface mount operational amplifiers withstand brief heating to 300 °C. We used SMD291AX soldering paste for SMD components and multicore welding wire (Sn63/Pb37) for other soldering. Soldering paste was applied by dot dispensing and we used fine soldering iron set to 250 °C maximum. Usually, only few seconds of heating are required to fix the components. After assembly, PCBs were soaked in isopropyl alcohol for 10 min, then extensively brushed using painting brush, rinsed again in clean isopropyl alcohol, wiped and dried.

### 4. Engineering details drawings of the enclosure cases.

The required modifications are shown in red.

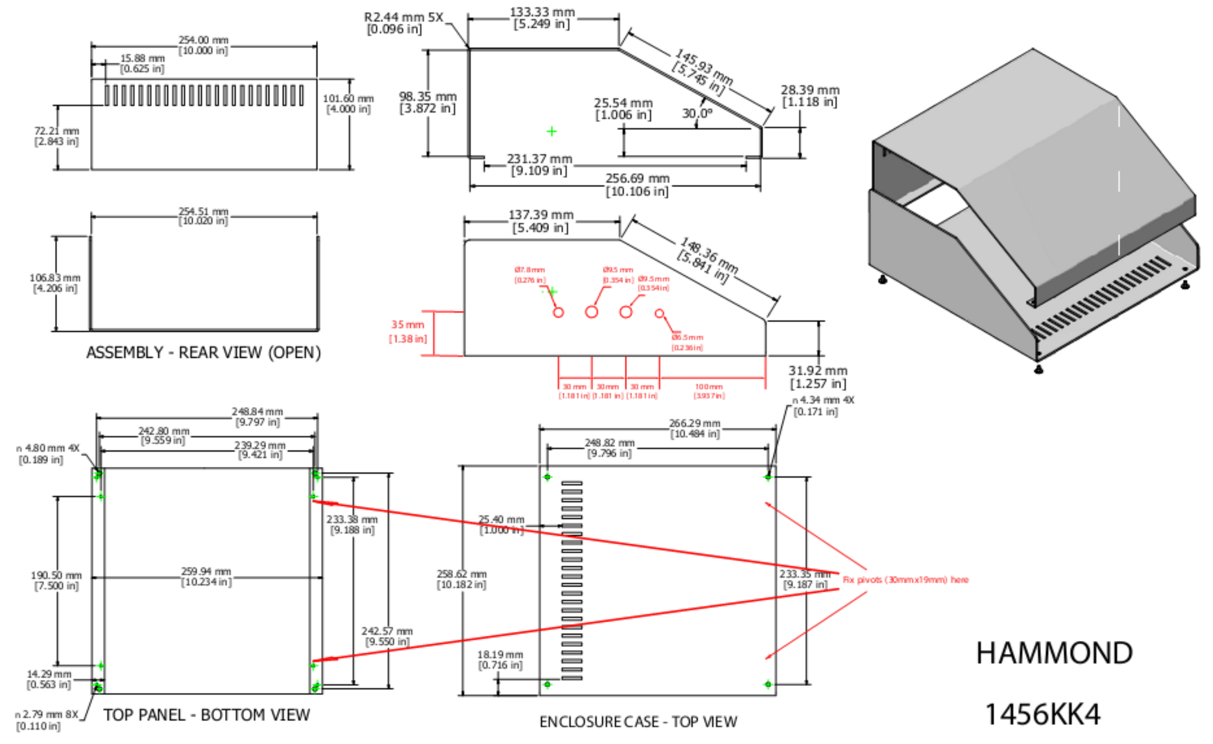


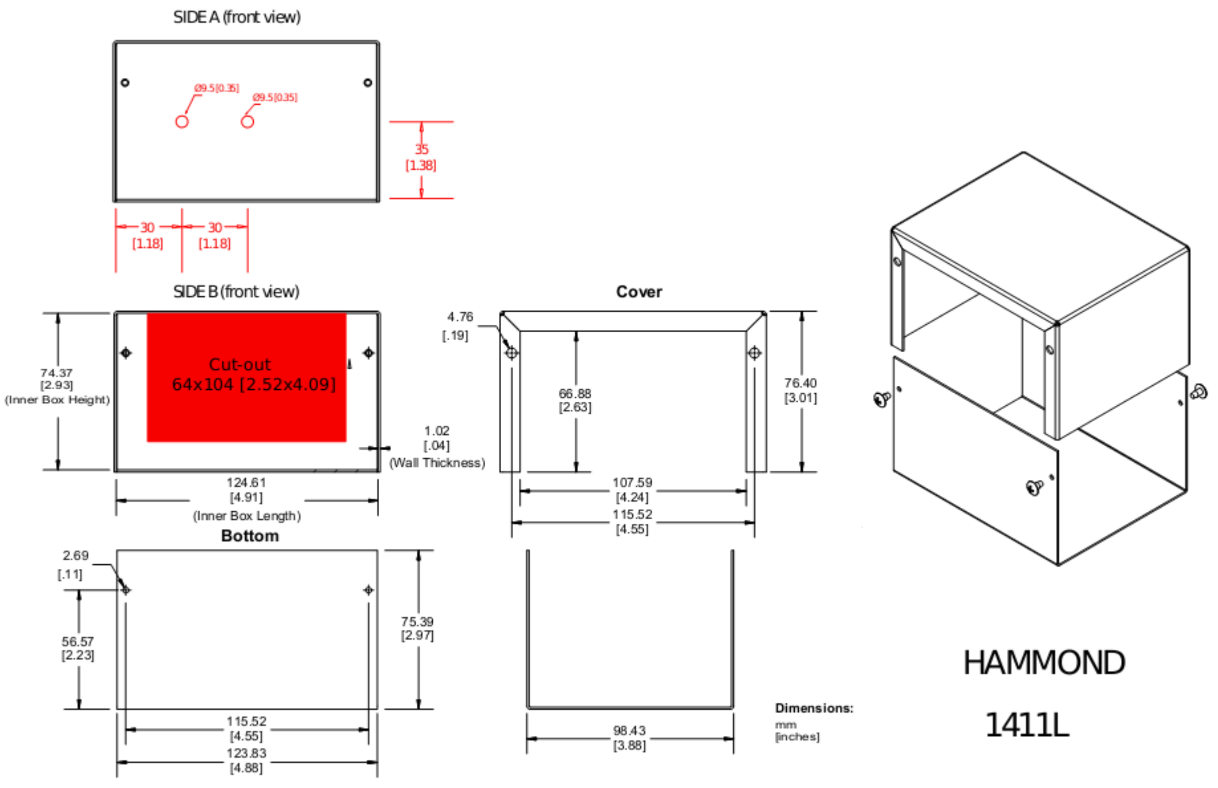

### 5. Theoretical and practical design of a high-speed low-noise headstage.

In this part, we revise several high-speed and low-noise approaches in voltage-clamp method in order to design a wideband open-source picoampere amplifier. Several great publications provide an in-depth overview of the electronic designs of a voltage-clamp amplifier for single-channel recording (Sigworth 1983, Alvarez 1986; Auerbach and Sachs 1985, Hanke and Schlue 1993, Sherman-Gold 1993). This supplementary information will consider the aspects of the electronic circuitry involved in frequency-response correction of the current-to-voltage converter, a central unit of voltage-clamp amplifier. In general, such frequency-response correction is necessary because classical (resistive) current-to-voltage (*I-V*) conversion in voltage-clamp amplifiers is carried out by operational amplifier with gigaOhm resistor ($R_f$) mounted in his negative feedback loop (Fig 1).

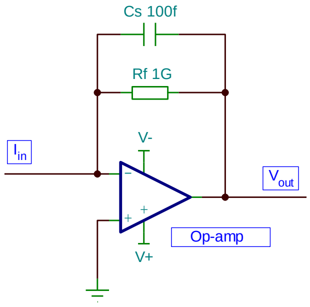


Figure 1. I-to-V converter

Unfortunately, all commercial resistors have small stray capacitance ($C_s$) distributed in parallel in the order of hundreds of femtoFarads. Then, even in case of high-speed operational amplifier, his transfer function is limited by a roll-off generated by low-pass $R_fC_s$ circuit with a corner frequency (at -3 dB) determined as $f_c=1/(2\pi R_fC_s)$. This suggests a bandwidth of 530 Hz for a 300 fF stray capacitor in parallel with 1 GΩ resistor, which obviously is not convenient for high-speed single-channel recordings. Accordingly, a technique that decreases stray capacitance in feedback resistors such as “field-shunt” is one of the ways to extend the bandwidth. It consists in shielding the feedback stray capacitance using a ground trace around the feedback resistor near the output side. This way, parasitic capacitance decreases below 100 fF, so the bandwidth extends by a factor of 3 or even better, yet this improvement is still not sufficient for wide-band voltage clamping. The more impactful methods of frequency response correction of transimpedance amplifiers (i.e. *I-V* convertors) are divided into three groups: 1) a “bootstrapping technique" that uses sophisticated circuitry eliminating the high-frequency current loss through the power supply capacitance; 2) introduction of compensatory RC T-network in the feedback loop of transimpedance amplifier; and 3) cascading *I-V* convertor with a high-pass shelving filter. The two latter techniques use discrete components, require minimal tuning and are relatively easy to implement, so only they are reviewed in the present work.

**Methods**

Analog circuit schematics design and virtual performance testing was accomplished using TINA-TI™ v9, a free version of SPICE-based tool developed by Texas Instruments and DesignSoft. Conventional DC, transient, frequency domain and noise analysis have been employed. Lipid bilayer membrane was modeled using a variable capacitor ($C_m$) in parallel with 100 GΩ resistor ($R_m$). Ion channel opening in transient analysis was simulated by the commutation of 100 GΩ resistor ($R_i$) in parallel with $R_m$ using time-controlled switch that has infinite $R_{off}$ and $R_{on}$=1Ω (Figure 2).

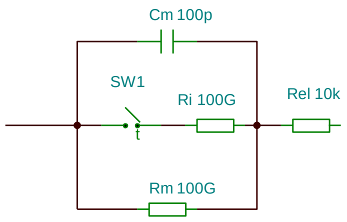


Figure 2. Model membrane circuit

During transient analysis, constant DC voltage was applied to one side of this $C_m$//$R_m$//$R_i$ circuit via 10 kΩ resistor representing total resistance of a pair of Ag/AgCl electrodes ($R_{el}$). The output membrane current was fed to the input of *I-V* converter. Accordingly, the side of voltage application made *cis*-compartment, while the opposite side was connected to the virtual ground and made *trans*-compartment. Circuits were powered by ±5 V bypassed by 10 µF and 10 nF capacitors. In noise analysis, total $V_{rms}$ noise was calculated. Transient analysis was performed at 0.1 µs temporal resolution. The bandwidth (BW) of the complete setup in the presence of different membrane capacitances was estimated from the measurement of the rise time ($\tau_{10\text{-}90}$) of the simulated transient output voltage during commutation of 100 GΩ resistor using following equation derived from the voltage relaxation equation of RC circuits (Sherman-Gold, 1993):

$$\mathbf{BW=0.349/\tau_{10\text{-}90}}$$

### Prototype testing

Eight-pole Bessel filter (LPF8) was used for signal conditioning prior to analysis using digital storage oscilloscope with adjustable sample rate (ISDS205B, Harbin Instrustar Electronic Technology, China). The bandwidth of the complete setup was estimated from the analysis of the output rectangular signal obtained by the application of triangular voltage waveform (10 V/s, 500 Hz) from the signal generator (GW Instek SFG2120) to the RC test-circuit composed of 10 kΩ resistor in series with parallel 1 GΩ resistor and 100 pF, 10 pF or 1 pF capacitor. The rise time ($\tau_{10\text{-}90}$) of 50 sweeps was averaged and presented as Mean±SD. The corresponding bandwidth was calculated using the equation above. According to Nyquist sampling theory, output $V_{rms}$ noise at a given bandwidth was measured using the sampling rate corresponding to 10-fold the bandwidth temporal resolution.

### T-network

Compensatory RC T-network for transfer function extension has been proposed in several old designs of transimpedance amplifiers (Goto and Ishikawa, 1979; Michel et al, 1992; Hanke and Schlue 1993) and was used in recent works in more complicated form (Howard 1999; Seidl et al, 2003; Giusi et al, 2014; Kim and Kim, 2018). Figure 3 shows a basic circuit with the T-network in the feedback loop.

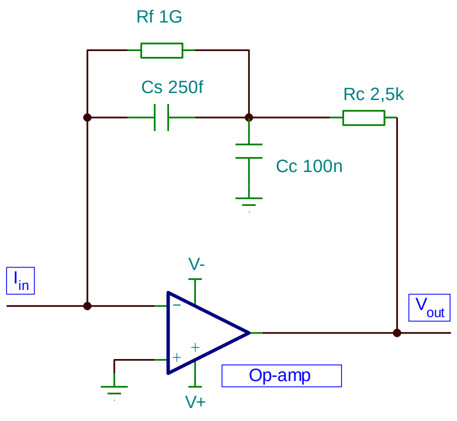


Figure 3. T-network compensatory circuit

At low signal frequencies, capacitances $C_s$ and $C_c$ are open circuits, so the DC voltage drop at the output of amplifier is determined by the sum of two resistors, $R_f$ and $R_c$. If this T-network is designed to respect the proportion of $R_f*C_s=R_c*C_c$, then starting at the low-pass corner frequency determined by $R_f$ and $C_s$, the network provides low impedance path to the ground, thus increasing the voltage drop at $R_c$. Accordingly, this keeps the signal magnitude at the level of DC signal which extends the transfer function of the amplifier. In order to avoid a creation of interfering low-pass pole at the $R_c$ due to its own parasitic capacitance, this resistor should be chosen as low as possible (usually in kΩ range). The advantage of this compensation technique is that it requires only one adjustable component, $R_c$, while $C_c$ can be fixed. Unfortunately, this design was susceptible to several drawbacks in our computer circuit analysis using transient response. First, it presented with significant ringing (overshot). This problem could be eventually solved using low-pass filter at the expense of losing some bandwidth. Second, the bandwidth of the amplifier was strongly dependent on the input (membrane) capacitance and dropped below 10 kHz at $C_m$=100 pF.

This technique was tested in a prototype. Most operational amplifiers tested were unstable in this circuit and required significant compensation capacitance in the order of several pF. Bandwidth and noise characteristics of the only one prototype with a minimal compensation of 3 pF is presented in Table 1. As can be seen, the actual maximal achievable bandwidth in the presence of 100 pF input capacitance is lower than 6 kHz. Moreover, amplifier noise at low frequencies was too high. In conclusion, this design is not the option to work with conventional lipid bilayers that have capacitances in the order of hundreds of pF.

**High-pass shelving filter**

A high-pass shelving filter used in electrophysiology increases the magnitude of signal (20 dB/decade) at frequencies above the frequency of *I-V* converter roll-off and keeps DC signal at unity gain (0 dB). Shelving filter is a first order filter that implements operational amplifiers over inverting or non-inverting topologies. Since the low-pass roll-off of *I-V* converter is also first order at 20 dB/decade, high-pass shelving filter perfectly compensates frequency response and keeps it flat at 0 dB. Accordingly, the output roll-off of the whole amplifier (i.e. including *I-V*-converter) is of the second order (40 dB/decade). Electrophysiology electronics borrowed shelving filters from analog audio equalizer designs.

For a long time before their application in electrophysiology, high- and low-pass shelving filters were cascaded in audio equalizers to provide a control over treble and bass frequencies, respectively. Accordingly, several high-pass shelving filters can be combined in series to extend frequency response of *I-V* converter to the limit determined only by the speed of operational amplifier used.

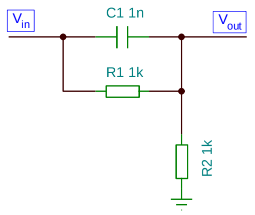


Figure 4. Passive high-pass shelving filter

*Non-inverting high-pass filter*

Non-inverting high-pass filter obtained the name “high-frequency booster” in the design of Hamill et al (1981). It is still used with some modification in resistive feedback headstage of Axopatch 200B amplifier, the benchmark for single-channel recordings. This booster circuit has been reviewed and explained in details in subsequent publications (Sigworth 1983, Alvarez 1986; Auerbach and Sachs 1985). The schematic of the filter is shown in Figure 5. In simulations, this design provides overall bandwidths in excess of 100 kHz, which is only slightly dependent on membrane input capacitance.

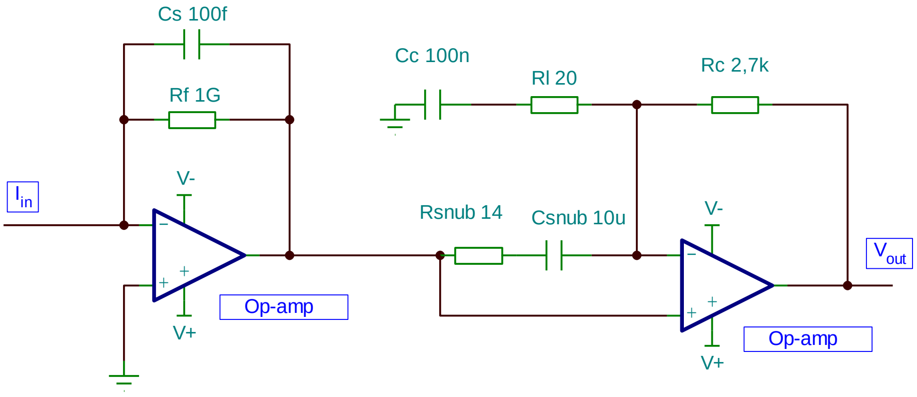


Figure 5. IV converter and non-inverting shelf filter with a snubber

We have built a number of prototypes based on this design to test their performance with available low-noise, low-offset and high-speed operational amplifiers. This design had a number of advantages compared to T-network; it introduces limiting potentiometer $R_l$ that serves to eliminate the overshot/ringing of the amplifier. Moreover, any desired bandwidth can be trimmed using this potentiometer. There are only two variable components in the filter, $R_c$ and $R_l$. Another advantage of non-inverting high-pass filter is that it provides output signal in phase with *I-V* converter, thus, it remains inverted; accordingly, it can be followed by the inverting gain voltage amplifier in order to obtain overall output voltage in phase with the input signal. However, despite of all our efforts to keep the amplifier stable at the broad bandwidth in the presence of high input capacitance ($C_m$=100 pF), most prototype amplifiers started to oscillate beyond 50 kHz even with a RC snubber compensation circuit (Figure 5). Table 1 shows few prototype examples with the bandwidth over 50 kHz, but, as can be seen, we have never reached our goal of 100 kHz. Apparently, the bandwidth of 50 kHz corresponds to the natural stability threshold of shelving filters in non-inverting configuration (or the stability of the overall cascade) in the presence of high input capacitance. This may probably explain why the frequency response of the resistive feedback headstage of Axopatch 200B is trimmed to 50 kHz. To confirm this, we have measured the rise time of Axopatch 200B with 100 pF membrane capacitance, which was equal to 7.75±0.65 μs that corresponds to the effective bandwidth of 45 kHz.

An interesting design of high-pass filter that gives the output signal in phase with *I-V* converter and thus may be considered as non-inverting high-pass filter has been presented by Kim and Koo (2005). The proposed topology implements inverting input stage with embedded inverting gain and is shown in simplified form in Figure 6.

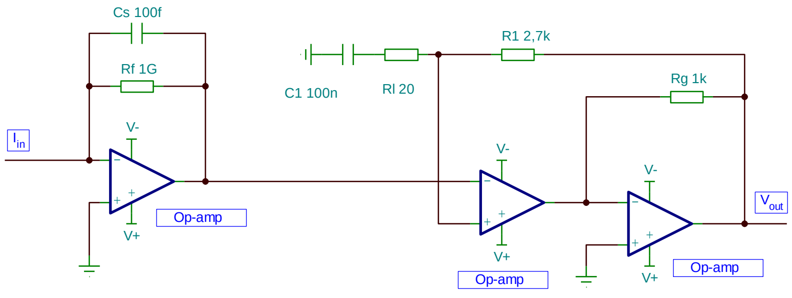


Figure 6. IV converter and non-inverting shelf filter with embedded gain.

In simulations, this design has several advantages compared to classical non-inverting high-pass shelving filter. First, the embedded gain improved stability at high frequencies, so we were able to obtain several designs with overall bandwidth of 100 kHz (Table 1). However, not any operational amplifier was stable in this design. Second, the embedded gain increased the sharpness of the output roll-off by one order (to 60 dB/decade), which significantly improved noise characteristics at high frequencies in simulations. Regrettably, this design showed no noise improvement in real prototypes at high frequencies (Table 1), which suggests its high sensitivity to parasitic electronics coupling.

*Inverting high-pass filter*

The use of inverting high-pass filter in electrophysiological voltage-clamp amplifier has been first proposed by Fleischmann and colleagues in 1980 (Fleschmann et al, 1980). Afterwards, it has been implemented in Dagan3900 amplifier (Dagan manual) and was used in several more recent designs with some modifications (Carlà et al 2004, Paul and Marsala 2006, Ciofy et al 2006, Ferrari and Sampietro2007, Hartel et al 2018).

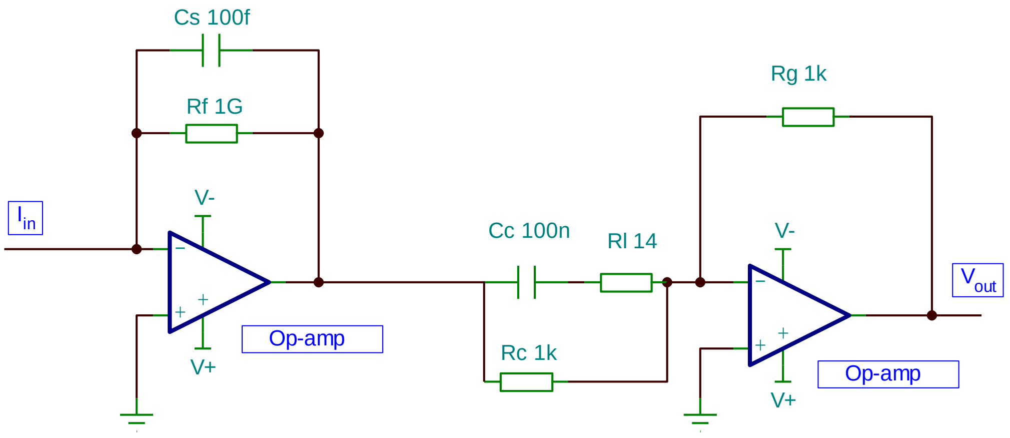


Figure 7. IV converter and inverting shelf filter

The basic schematic of the conventional inverting high-pass shelving filter is shown in Figure 7. In 1989, Strickholm proposed to separate DC and AC signals in his hybrid voltage-clamp amplifier, where the inverting shelving filter was applied only to the AC path of the signal (Strickholm, 1989). However, in our computer simulations, this approach provided no advantage in terms of frequency response correction and noise compared to conventional design. In addition, it has more components and requires more tuning, so it has not been prototyped in our study. As can be seen in Figure 7, conventional inverting high-pass shelving filter design has three variable components, $R_c$, $R_l$ and $R_g$. While the two former potentiometers together with compensatory capacitance $C_c$ have the same functions as in non-inverting shelving filter, gain potentiometer $R_g$ sets the output DC amplitude irrespective of frequency correction. This is advantageous because of the following reason: usually it is difficult to find feedback resistor with the exact 1 GΩ value (usual tolerance is ±5%), so the accurate transimpedance gain can be adjusted using $R_g$ potentiometer. This is impossible in the non-inverting topology. Computer simulations suggested that limiting potentiometer ($R_l$) in basic schematics is not that efficient in preventing amplifier overshot and ringing. The alternative method of ringing limitation is to form a low-pass pole in the feedback loop of shelving amplifier using a capacitor. However, trimming capacitors are usually bulky and not practical. Instead, the same work can be performed by so-called snubber RC network placed between the two inputs of shelving amplifier. In order to snub effectively only high frequency ringing, a high quality (low ESR) and large value ($C_l$ =10-100 μF) ceramic capacitor should be connected in series with the low-value potentiometer $R_l$. The updated design is presented in Figure 8.

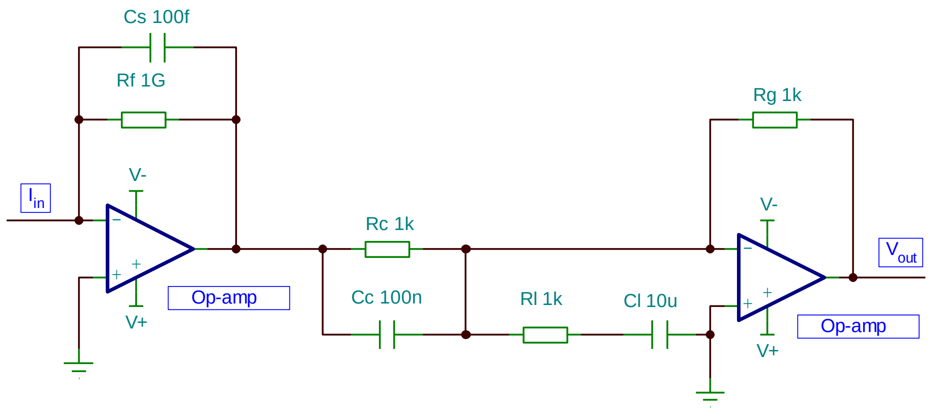


Figure 8. IV converter and inverting shelf filter with a snubber.

The performance of a number of prototypes is shown in Table 1. Clearly, many prototypes provide stable bandwidths in excess of 100 kHz in the presence of 100 pF input capacitance. Noise characteristics are shown in Table 1 and they are comparable to that of the state of the art amplifier Axopatch200B.

Finally, by the analogy to the design of Kim and Koo (2005) we have developed and investigated an inverting shelf filter with embedded gain. This design is presented in Figure 9.

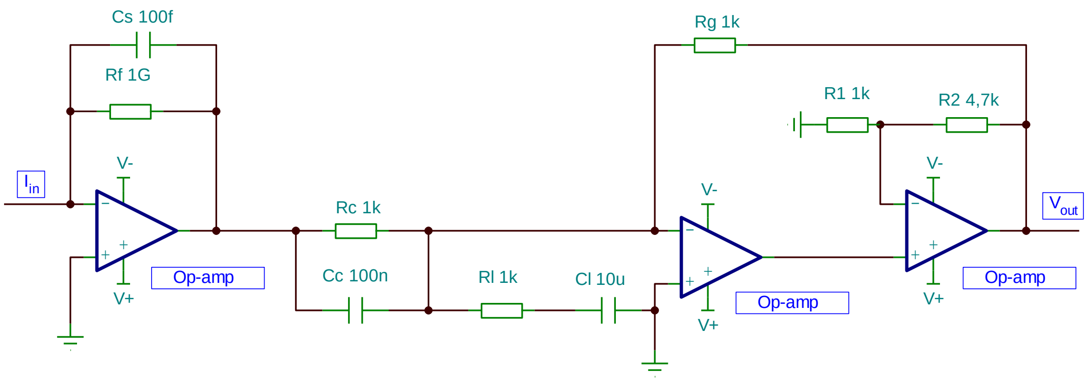


Figure 9. IV converter and inverting shelf filter with a snubber and with embedded gain.

Although this design performed very well in computer simulations and provided the bandwidths in excess of 500 kHz in some instances, it was prone to oscillations in prototypes. It required very fine tuning that depended on the variations of the input capacitance. It seems that this design is extremely sensitive to the parasitic electronic components present on PCBs.

Overall, our prototyping efforts and electronic tests suggest that IV converter with inverting shelf filter stabilized by a snubber network is the most stable design. It is also evident that the designs using ADA4817-1 operational amplifier provide both required bandwidth and acceptable noise characteristics (Figure 8, Table 1). This updated design and ADA4817-1 have been chosen for the realization of the OpenPicoAmp-100k amplifier.

### Table 1. Amplifiers bandwidths and noise

| Resistive headstage AXOPATCH 200B : $pA_{RMS}$ noise with open input | | | | | | Resistive headstage AXOPATCH 200B : $pA_{RMS}$ noise with 100 pF/10 kΩ | | | | | |
|---|---|---|---|---|---|---|---|---|---|---|---|
| 100 kHz | 40 kHz | 20 kHz | 10 kHz | 1 kHz | 0,1 kHz | 100 kHz | 40 kHz | 20 kHz | 10 kHz | 1 kHz | 0,1 kHz |
| 51.7 | 9.8 | 4.8 | 2.51 | 0.52 | 0.071 | 101 | 39.5 | 18.5 | 7.65 | 0.67 | 0.095 |

**T-NETWORK**

$R_f$=1G ;$C_f$=3pF ;$R_c$=10k(variable) ;$C_c$=470nF

| **Amplifier components** | | | **$T_{10-90}$,** | **BW** | **$pA_{RMS}$ noise with open input** | | | | | | | **$pA_{RMS}$ noise with 100 pF/10 kΩ** | | | | | | |
|---|---|---|---|---|---|---|---|---|---|---|---|---|---|---|---|---|---|---|
| Opa1 | | $R_c$ | µs | **kHz** | BP | 100 kHz | 40 kHz | 20 kHz | 10 kHz | 1 kHz | 0,1 kHz | BP | 100 kHz | 40 kHz | 20 kHz | 10 kHz | 1 kHz | 0,1 kHz |
| LM6211 | | 6k | 62±1 | 5.6 | | | | | 3.28 | 0.77 | 0.095 | | | | | 3.71 | 1.05 | 0.135 |

**NON-INVERTING SHELVING FILTER WITH SNUBBER**

$R_f$=1G ;$C_f$=270fF ;$R_c$=20k(variable) ;$C_c$=109 nF ; $R_l$=1k

| **Amplifier components** | | | $T_{10-90}$, | **BW** | **$pA_{RMS}$ noise with open input** | | | | | | | **$pA_{RMS}$ noise with 100 pF/10 kΩ** | | | | | | |
|---|---|---|---|---|---|---|---|---|---|---|---|---|---|---|---|---|---|---|
| Opa1 | Opa2 | $R_c/R_{lim}$ | µs | **kHz** | BP | 100 kHz | 40 kHz | 20 kHz | 10 kHz | 1 kHz | 0,1 kHz | BP | 100 kHz | 40 kHz | 20 kHz | 10 kHz | 1 kHz | 0,1 kHz |
| ADA4817 | LT1028 | 13870/39 | 4.87±0.47 | 71.9 | | 9.4 | 3.0 | 1.61 | 0.90 | 0.16 | 0.051 | | 150 | 58 | 19.9 | 7.46 | 0.39 | 0.063 |
| AD8067 | LT1028 | 3815/46 | 4.76±0.66 | 73.5 | | 94 | 16.5 | 7.75 | 3.35 | 0.27 | 0.065 | | 246 | 80 | 26.9 | 11.12 | 0.46 | 0.072 |
| AD8033 | LT1028 | 2860/39 | 4.40±0.50 | 79.5 | | 84.5 | 10.5 | 4.24 | 1.75 | 0.17 | 0.055 | | 270 | 100 | 29.6 | 10.91 | 0.45 | 0.069 |
| ADA4637 | LT1028 | 3648/39 | 3.96±0.56 | 88.4 | | 56 | 8.4 | 3.87 | 1.81 | 0.21 | 0.058 | | 249 | 83 | 21.5 | 7.89 | 0.35 | 0.061 |

**NON-INVERTING SHELVING FILTER WITH EMBEDDED GAIN**

$R_f$=1G ;$C_f$~270f ;$R_c$=10k(variable) ;$C_c$=116nF ; $R_{lim}$=500(variable) ; $R_g$=1k ; field shunt applied

| **Amplifier components** | | | **$T_{10\text{-}90}$,** | **BW** | **$pA_{RMS}$ noise with open input** | | | | | | | **$pA_{RMS}$ noise with 100 pF/10 kΩ** | | | | | | |
|---|---|---|---|---|---|---|---|---|---|---|---|---|---|---|---|---|---|---|
| Opa1 | Opa2 | $R_c/R_{lim}$ | µs | **kHz** | BP | 100 kHz | 40 kHz | 20 kHz | 10 kHz | 1 kHz | 0,1 kHz | BP | 100 kHz | 40 kHz | 20 kHz | 10 kHz | 1 kHz | 0,1 kHz |
| AD8067 | OPA2227 | 3280/28 | 3.24±0.67 | 108 | 69 | 31 | 14.7 | 6.8 | 2.92 | 0.24 | 0.060 | 302 | 157 | 63.7 | 18.7 | 6.95 | 0.35 | 0.071 |
| AD8067 | TL072CN | 4058/38 | 4.50±0.67 | 78 | 63 | 28 | 12.6 | 6.1 | 2.42 | 0.21 | 0.059 | 160 | 110 | 55.7 | 18.1 | 6.95 | 0.35 | 0.071 |

**INVERTING SHELVING FILTER WITH SNUBBER**

$R_f$=1G ;$C_f$~270f ; $R_g$=$R_c$=20k(variable) ;$C_c$=116nF ; $C_{lim}$=10 µF, $R_{lim}$=500(variable) ; field shunt applied

| **Amplifier components** | | | $T_{10\text{-}90}$, | BW | **$pA_{RMS}$ noise with open input** | | | | | | | **$pA_{RMS}$ noise with 100 pF/10 kΩ** | | | | | | |
|---|---|---|---|---|---|---|---|---|---|---|---|---|---|---|---|---|---|---|
| Opa1 | Opa2 | $R_g$=$R_c/R_{lim}$ | µs | **kHz** | BP | | 40 kHz | 20 kHz | 10 kHz | 1 kHz | 0,1 kHz | BP | | 40 kHz | 20 kHz | 10 kHz | 1 kHz | 0,1 kHz |
| AD8067 | TLC070 | 3920/43 | 2.29±0.84 | 152 | 143 | | 38.5 | 7.81 | 2.79 | 0.21 | 0.051 | 615 | | 68 | 22.6 | 7.57 | 0.80 | 0.195 |
| ADA4817 | AD8067 | 9890/3.1 | 2.36±0.59 | 148 | 26.7 | | 4.72 | 2.79 | 1.61 | 0.17 | 0.055 | 375 | | 49.1 | 18.1 | 6.87 | 0.99 | 0.350 |
| AD8067 | ADA4622 | 3847/39 | 2.54±0.54 | 137 | | | | | | | | 395 | | 83 | 24.4 | 7.95 | 0.55 | 0.195 |
| AD8067 | LT1007 | 3802/41 | 3.35±0.33 | 104 | 92 | | 30 | 7.41 | 2.58 | 0.20 | 0.051 | 365 | | 61 | 20.0 | 7.11 | 0.65 | 0.147 |
| AD8067 | OPA227 | 3781/40 | 3.44±1.00 | 101 | 79 | | 28 | 8.05 | 2.79 | 0.21 | 0.053 | 382 | | 62 | 22.5 | 8.10 | 0.77 | 0.195 |
| ADA4817 | ADA4637 | 9240/7.9 | 3.58±0.55 | 98 | 12.1 | | 2.95 | 1.59 | 0.95 | 0.14 | 0.049 | 187 | | 44.3 | 16.5 | 6.05 | 0.31 | 0.059 |
| ADA4817 | LT1115 | 14481/17 | 4.04±0.92 | 86 | 12.7 | | 4.50 | 1.74 | 0.89 | 0.16 | 0.051 | 350 | | 57.5 | 21.5 | 7.55 | 0.75 | 0.151 |
| ADA4817 | ADA4625 | 14476/25 | 6.65±0.47 | 53 | 5.65 | | 2.83 | 1.52 | 0.82 | 0.14 | 0.052 | 94 | | 40.5 | 18.9 | 7.12 | 0.36 | 0.059 |
| ADA4817 | LT1007 | 12360/79 | 11.51±0.49 | 30 | 2.79 | | 1.86 | 1.28 | 0.78 | 0.14 | 0.052 | 41.5 | | 25.5 | 15.1 | 6.35 | 0.35 | 0.060 |
